# Lipid Packing Defects are Necessary and Sufficient for Membrane Binding of α-Synuclein

**DOI:** 10.1101/2024.11.14.623669

**Authors:** David H. Johnson, Orianna H. Kou, John M. White, Stephanie Y. Ramirez, Antonis Margaritakis, Peter J. Chung, Vance W. Jaeger, Wade F. Zeno

**Affiliations:** Mork Family Department of Chemical Engineering and Materials Science, University of Southern California, Los Angeles, CA, 90089, United States; Department of Physics and Astronomy, University of Southern California, Los Angeles, California, 90089, United States; Department of Chemical Engineering, University of Louisville, Ernst Hall, Room 312, 216 Eastern Parkway, Louisville, Kentucky 40292, United States; Department of Biological Sciences, University of Southern California, Los Angeles, 90089, United States; Department of Chemistry, University of Southern California, Los Angeles, California, 90089, United States; Alfred E. Mann Department of Biomedical Engineering, University of Southern California, Los Angeles, California, 90089, United States

## Abstract

α-Synuclein (αSyn), an intrinsically disordered protein implicated in Parkinson’s disease, is thought to initiate aggregation by binding to cellular membranes. Previous studies suggest that anionic lipids are necessary for this binding. However, these studies largely focused on unmodified αSyn, while physiological αSyn is N-terminally acetylated (NTA). Our work challenges the long-standing paradigm that anionic lipids are necessary for αSyn binding by demonstrating that NTA diminishes αSyn’s reliance on anionic membrane charge, revealing that membrane packing defects (i.e., interfacial hydrophobicity) alone can drive membrane binding. Using fluorescence microscopy and circular dichroism spectroscopy, we monitored the binding of NTA-αSyn to membrane vesicles with different lipid compositions. Phosphatidylcholine and phosphatidylserine concentrations were varied to control surface charge, while phospholipid tail unsaturation and methylation were varied to modulate lipid packing. We also formulated cholesterol-containing membranes that mimicked the lipid composition of synaptic vesicles. In these membranes, all-atom molecular dynamics simulations were used to visualize and quantify membrane packing defects. Our results demonstrate that membrane packing defects are necessary for NTA-αSyn binding and that defect-rich membranes are sufficient for NTA-αSyn binding regardless of membrane charge. These findings provide a molecular mechanism by which lipid structural properties, such as poly-unsaturation, can regulate αSyn binding to physiological membranes.

## Introduction

α-Synuclein (αSyn) is a membrane-binding intrinsically disordered protein found primarily in the brain, where it localizes to synaptic terminals of neurons ^1^. Its notoriety originates from its implication and pathogenic function in neurodegenerative diseases, most notably in Parkinson’s Disease (PD), through its inclusion in Lewy bodies ^2–4^. However, in recent years, the intimate relationship between α-synuclein’s physiological function and pathology has become more apparent ^5^. This revelation has led to a need to better understand the fundamental interactions that dictate αSyn’s function, specifically in its interaction with lipid bilayers ^6^. For example, αSyn is known to assist in the trafficking of synaptic vesicles ^7–9^, but recent research has shown that membrane lipid composition, specifically, affects the role of αSyn in vesicle clustering and docking during trafficking ^10–12^. Additionally, it is known that monomeric αSyn interactions with lipid membranes influence or even induce ^13^ fibrilization and aggregation of αSyn ^14–16^. These insights bring attention to the overall importance of how αSyn-membrane interactions dictate αSyn function and pathology.

Over the last twenty years, many studies have described the binding behavior of αSyn to a variety of vesicle compositions to better understand its purpose and function ^10,17–29^. Upon membrane association, this 140 amino-acid protein transitions from a completely structureless state in solution to a partially folded structure on the membrane surface, where the N-terminal domain consisting of approximately 100 residues adopts an α-helical conformation ^21,30^. One primary observation from these studies is the critical importance of anionic phospholipids for αSyn binding ^19–22,24,28^. Previous work has also demonstrated that αSyn binding is influenced by lipid packing defects, which can be defined as transient surface gaps between lipid headgroups where the underlying hydrophobic hydrocarbon tails are exposed to solvent ^31^. It is important to note that a lipid packing defect, as defined here, is a term to quantify exposed hydrophobic area of lipid bilayers due to non-ideal packing, highlighting the importance of hydrophobic interactions for aSyn binding. These packing defects have been shown to enhance membrane binding of wild type αSyn (WT-αSyn) ^26,27^, especially when phospholipids containing methyl-branched tails ^17^ or single tails ^12,32^ were used. While there is a substantial body of work collectively describing the effects of familial PD point mutations on αSyn- membrane interactions ^20,33,34^, fundamental studies on the effects of post-translational modifications (PTMs) have only been more recently considered as many αSyn PTMs are found to be implicated in aggregation, formation of Lewy bodies, and neurodegeneration ^35–37^. As a primary example, physiological αSyn exists with an N-terminal acetylation ^38–42^. Importantly, this PTM has been shown to modify αSyn aggregation kinetics ^43–45^ and affinity for lipid membranes ^46–49^ without modifying its structural properties when membrane-bound ^50^. This PTM has been overlooked in many previous binding studies, but it is important to consider when understanding αSyn’s function and pathology due to its physiological relevance and abundance ^38,39^. To date, existing literature comparing the binding behavior between WT-αSyn and N-terminally acetylated αSyn (NTA-αSyn) shows minimal difference in binding behavior to highly charged membranes containing 50 mol% of anionic phosphatidylserine lipids ^47^. This difference was more pronounced in moderately charged membranes containing 15-25% phosphatidylserine, where NTA-αSyn exhibited greater membrane binding than WT-αSyn ^46,49^. However, the effect of NTA on the binding of αSyn in the context of membrane defects remains to be studied, leaving a gap in understanding of how lipid packing defects impact the binding of NTA-αSyn in the context of membrane charge. By considering membrane charge, defect content, and N-terminal acetylation together, we hope to bring new insight into αSyn membrane interactions.

Here, we sought to investigate NTA-αSyn binding as a function of membrane charge and packing defect density by using a variety of lipid compositions in small unilamellar vesicles (SUVs). We first observed experimentally that NTA increased the binding of αSyn to both neutral and anionic membranes when compared to WT-αSyn but saw a disproportionate increase and preference for binding to defect-rich neutral membranes. This observation warranted a more rigorous study into the importance of packing defects for αSyn membrane binding. To quantify packing defect size and quantity, we established model systems using all-atom molecular dynamics simulations of lipid bilayers. The combination of experimental and simulation data revealed that as lipid packing defects increased across membrane compositions, NTA-αSyn binding also increased. Most notably, when anionic charge was added to membranes, NTA-αSyn binding did not increase unless the density of packing defects also increased. Therefore, our results suggest that charge alone is not enough to regulate NTA-αSyn binding and that membrane packing defects are necessary and sufficient for αSyn binding, differing from previous analyses that suggest anionic lipids are the primary promotor of membrane binding ^19,28^. Lastly, we show that membrane packing defects can be regulated in SUVs mimicking synaptic vesicle compositions by increasing the degree of unsaturation in phospholipid tails. Together, these observations reform the existing narrative about how αSyn binding is regulated for biological membranes and indicate that compositional considerations beyond phospholipid charge are necessary to fully encapsulate NTA-αSyn binding behavior.

## Results

### NTA-**α**Syn can exhibit preferential binding to neutral membranes over anionic membranes

To compare membrane compositional preferences of WT-αSyn and NTA-αSyn (**Figure 1A**), we utilized a previously developed, fluorescence microscopy-based tethered vesicle assay to monitor αSyn binding to SUVs ^51^ (**Figure 1B**). SUV diameters were determined through calibrations where dynamic light scattering (DLS) distributions and fluorescence intensity distributions were overlaid, while αSyn binding was quantified by performing single molecule imaging of fluorescently labeled proteins. Detailed explanations for these experimental procedures are provided in the Materials and Methods section. For these fluorescence microscopy experiments, αSyn was fluorescently labeled with ATTO-488 on residue 136, while SUV bilayers were doped with 0.5 mol% ATTO-647N labeled lipid. We used fluorescence microscopy and circular dichroism (CD) spectroscopy to verify that the inclusion of the fluorescent probe negligibly impacted αSyn’s ability to bind membranes (**Figure S1**). First, we compared the binding of αSyn to SUVs composed of 3:1 (mol:mol) dioleoylphophatidylcholine:dioleoylphosphatidylserine (DOPC:DOPS) and SUVs composed of diphytanoylphosphatidylcholine (DPhPC). The DOPC/DOPC SUVs represent anionic membrane surfaces with a 25% negative charge, while the DPhPC SUVs represent zwitterionic membrane surfaces with zero net charge. These two compositions were chosen because prior studies showed that tail methylation ^17^ and anionic headgroups ^19^ increased WT-αSyn binding. However, these properties have never been examined head-to-head, particularly for NTA-αSyn; therefore, we sought to observe αSyn binding preference to both membrane compositions.

**Figure 1.**
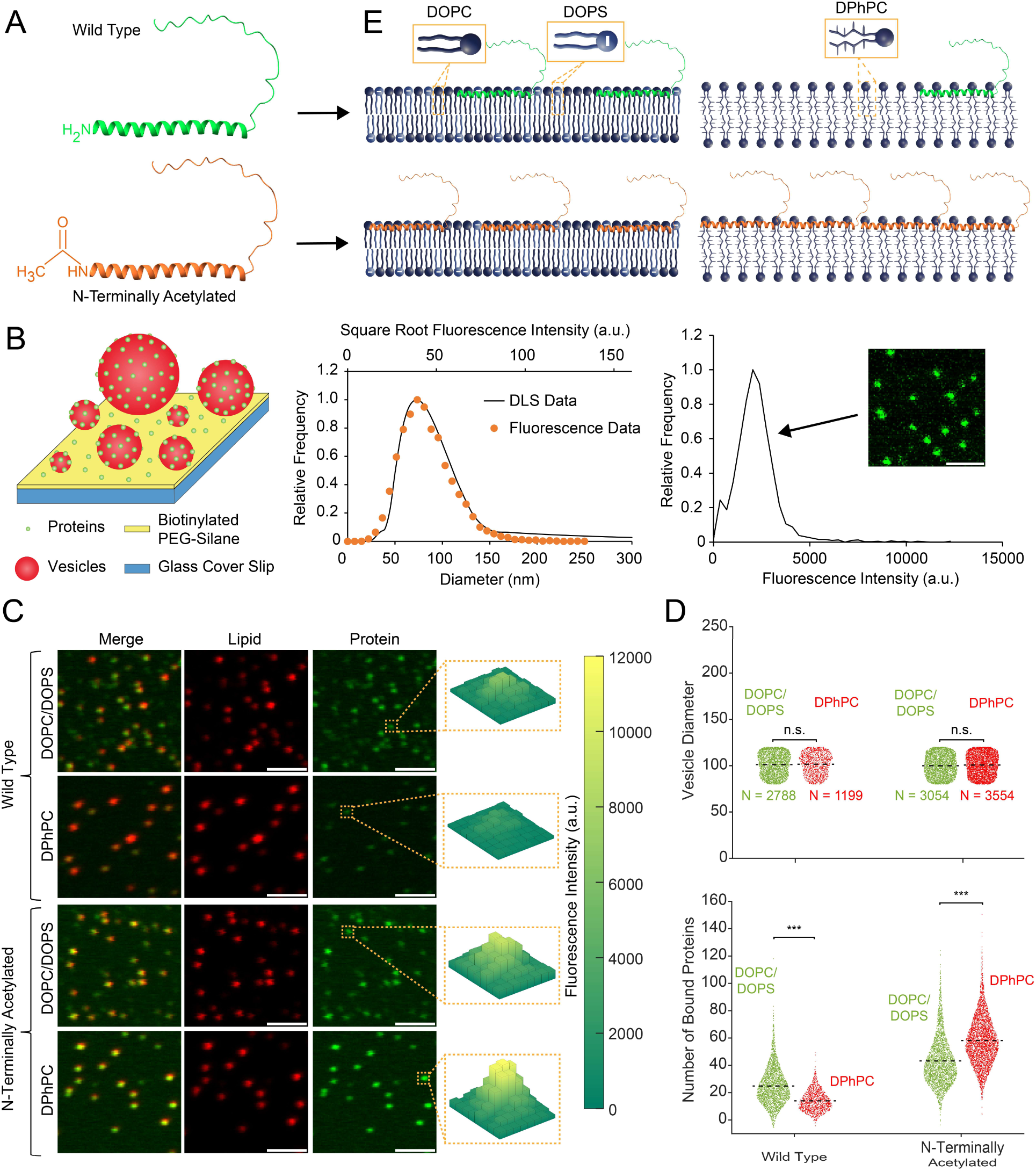
N-terminal acetylation of α-synuclein enhances overall membrane binding and sensitivity for lipid packing defects. (A) Structures of WT- and NTA-αSyn. (B) Schematic of the tethered vesicle assay (left) with a representative DLS/fluorescence intensity calibration plot for fluorescent SUVs (middle) and a representative single-molecule fluorescence intensity distribution for fluorescent αSyn (right). Scale bar represents 2 µM. (C) Lipid, protein, and merged fluorescence micrographs of tethered SUVs in the presence of 1 µM αSyn. Scale bars represent 2 µm. Yellow boxes highlight representative SUVs with bound αSyn and the corresponding three-dimensional intensity profiles in the αSyn fluorescence channels. (D) (top) The distribution of diameters for analyzed vesicles that were incubated with 1 µM αSyn and (bottom) the number of proteins bound to the individual SUVs from the top panel. (E) NTA-αSyn preferentially binds to neutral, defect-rich DPhPC membranes over anionic DOPC/DOPS membranes. DOPC/DOPS membranes contained a 3:1 (mol:mol) DOPC:DOPS. This schematic is a qualitative representation of αSyn-membrane association and is not intended to quantitatively reflect αSyn size and penetration depth relative to the membrane surface. The black dashed lines in D indicate the average values of the distributions. n.s. = not significant as determined by unpaired Student’s t-test. *** correspond to P < 0.001 as determined by Welch’s t-test (WT) and unpaired Student’s t- test (NTA).

In **Figure 1C**, αSyn binding was observed via fluorescence microscopy as the fluorescence signals from the protein channel were spatially colocalized with the fluorescence signals from the lipid channel. During our processing and analysis procedures for fluorescence images, we isolated SUV populations with an average diameter of 100 nm to control for membrane curvature effects (**Figure 1D**), as αSyn is known to sense membrane curvature ^52^. In **Figure 1C** WT-αSyn exhibited a higher fluorescence intensity on DOPC/DOPS membranes in comparison to DPhPC membranes, which corresponds to a higher degree of binding on DOPC/DOPS vesicles, as quantified in **Figure 1D**. Shown in **Figures 1C and 1D**, N-terminal acetylation increased the overall membrane affinity for both DOPC/DOPS and DPhPC membranes, which is consistent with previous observations ^46,47,49^. However, unlike WT-αSyn, NTA-αSyn displayed a binding preference for the neutral DPhPC SUVs. NTA resulted in a ∼2-fold increase for αSyn binding on DOPC/DOPS SUVs, while DPhPC SUVs elicited a ∼5-fold increase. These relative trends persisted on SUVs that were incubated with lower concentrations of WT-αSyn and NTA-αSyn (**Figure S2**). While there is a growing body of recent literature showing the importance of neutral membrane lipids for αSyn-membrane interactions ^11,12,32^, to the best of our knowledge, our results indicate the first reported instance of αSyn exhibiting a preference for neutral membranes over highly charged anionic membranes (**Figure 1E**). Importantly, these findings were obtained under conditions that controlled for membrane curvature (**Figure 1D**). Therefore, it is clear that the tail chemistry and, ultimately, lipid packing play a significant role in the membrane binding of NTA-αSyn. This fascinating result warranted further analysis to understand the mechanism and potential physiological implications for NTA-αSyn binding preferences to defect-rich membranes over charged membranes.

### NTA-**α**Syn membrane binding increases with higher packing defect content

Due to the various competing factors that control lipid packing, such as membrane curvature and membrane phase behavior, all lipids in **Figure 2** were analyzed at constant curvature (**Figure S3-S4**) in the liquid crystalline phase at room temperature (i.e., 21°C). **Figure 2A** provides structures for four phospholipids: dilauroylphosphatidylcholine (DLPC), DOPC, DOPS, and DPhPC. Collectively, these phospholipids represent the three types of tail chemistries analyzed in this work – dilauroyl (DL), dioleoyl (DO), and diphytanoyl (DPh), as well as two classes of headgroup – zwitterionic phosphatidylcholine (PC) and anionic phosphatidylserine (PS). The DL lipids were expected to exhibit the closest packing due to the complete saturation of their carbon chains, while the DO and DPh tails were expected to pack increasingly more loosely due to mono-unsaturation and methylation, respectively. PC and PS headgroups were chosen due to their physiological relevance throughout intracellular membranes ^53^. To quantify the relative quantity of packing defects in each membrane composition, we performed all-atom molecular dynamics simulations of lipid bilayers (**see Supplementary Information, Table S1-2**). **Figure 2B** shows selected simulation snapshots for pure DLPC and DPhPC membranes. Each snapshot represents a single frame from a simulation trajectory of 300 ns. In these snapshots, distinct differences in defect content for each composition can be noted, as the DPhPC membrane exhibited greater exposure of the tail groups, which are highlighted in yellow. Together, the combination of DL, DO, DPh tails and PC, PS headgroups yields 6 different phospholipids, 5 of which were examined in **Figure 2C**. Using the open-source software package PackMem ^54^, we quantified the average size (**Figure 2C, left)**, average number (**Figure 2C, center)**, and average membrane surface coverage (**Figure 2C, right)** of packing defects (refer to **Supplementary Information** for details of these calculations, **Figure S5**). For PC headgroups, the average size, average number, and surface coverage of defects were higher for DO tails than for DL tails. The inclusion of PS in 3:1 DLPC:DLPS membranes did not change the average size of packing defects compared to pure DLPC membranes, nor did it increase the frequency and surface coverage of packing defects. However, the inclusion of PS in 3:1 DOPC:DOPS membranes resulted in significant increases to defect size and quantity when compared to pure DOPC membranes. Interestingly, pure DPhPC membranes possessed substantially larger and more numerous defects than the other 4 systems examined.

**Figure 2.**
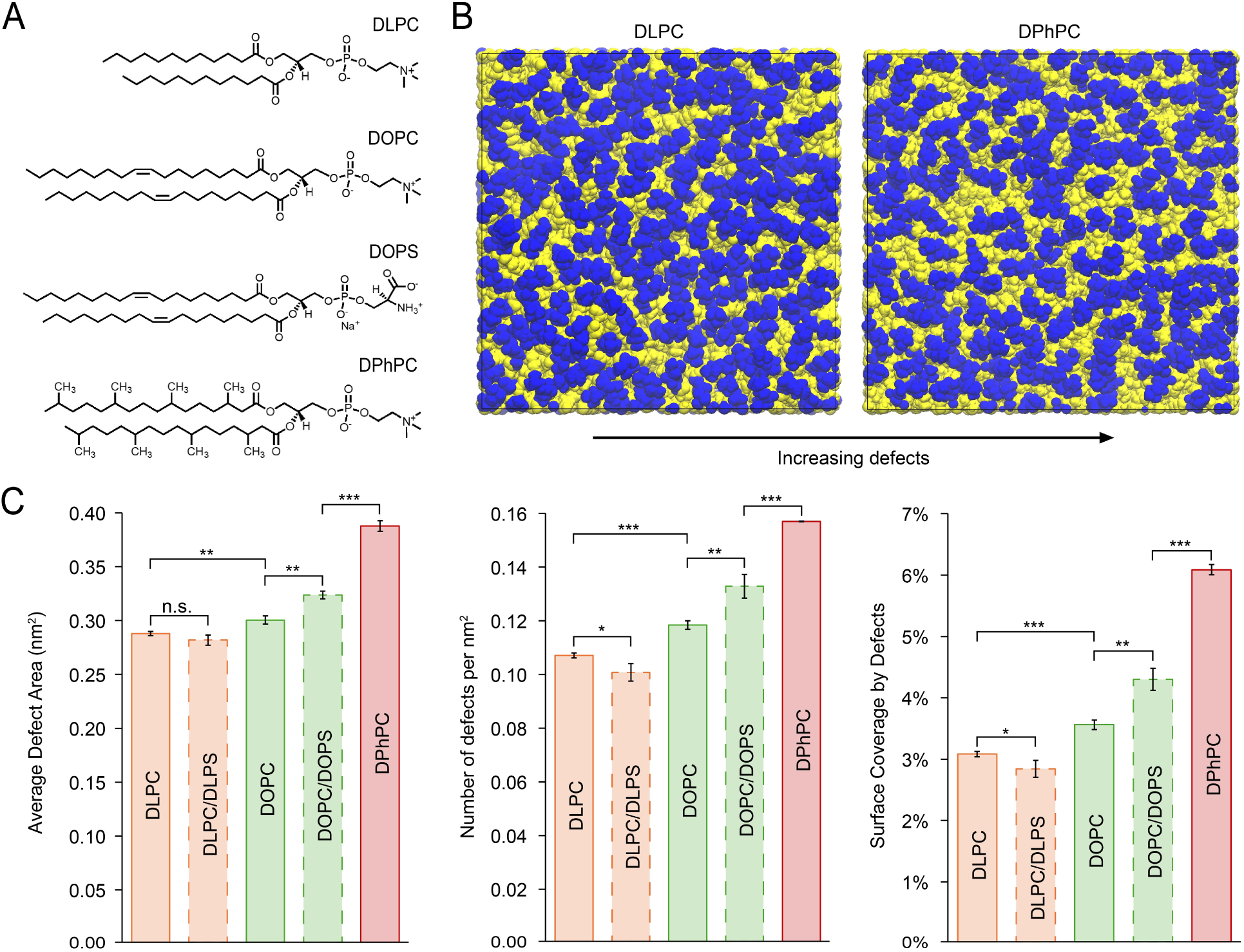
All-atom molecular dynamics simulations of lipid membranes allow for quantification of membrane packing defects. (A) Chemical structures for the primary phospholipid headgroups (PC, PS) and tail groups (DL, DO, DPh) used in the simulations. (B) Simulation snapshots of DLPC and DPhPC membranes showing increasing surface coverage of packing defects. Lipid tail groups are colored in yellow while lipid headgroups are colored blue for contrast. (C) The average area of a single packing defect in a simulation frame (left). The average number of packing defects per square nm in a simulation frame (center). The average surface coverage by membrane packing defects in a simulation frame (right). All PC/PS membranes contained a 3:1 PC:PS molar ratio. Error bars in (C) represent the standard deviation (SD) of 3 sets of simulations. *, **, and *** correspond to P < 0.05, P < 0.01, and P < 0.001, respectively, as determined by Student’s t-test with n = 3.

We next examined the binding preference of NTA-αSyn to SUVs as a function of lipid composition (**Figure 3**). Using the tethered vesicle assay to generate binding curves, we observed an increase in NTA-αSyn binding as the lipid composition was varied from pure DLPC to pure DOPC, then finally pure DPhPC (**Figure 3A, Figures S3-4)**. This increased binding was consistent with increased density of packing defects (**Figure 2C**). When 25% PS was included in DLPC/DLPS and DOPC/DOPS membranes, the resulting binding trends of the five lipid compositions examined mirrored the trends for defect coverage in **Figure 2C**, suggesting that the level of binding for NTA-αSyn may be proportional to the surface coverage of packing defects. Additionally, the results in **Figure 2C** and **Figure 3B** reveal that including 25% PS did not significantly increase the presence of defects nor the amount of NTA-αSyn binding for DL membranes. Conversely, 25% PS substantially increased the surface coverage of packing defects in DO membranes and the levels of NTA-αSyn binding. Nonetheless, neutral DPhPC membranes elicited the highest degree of NTA-αSyn binding for the 5 compositions examined so far.

**Figure 3.**
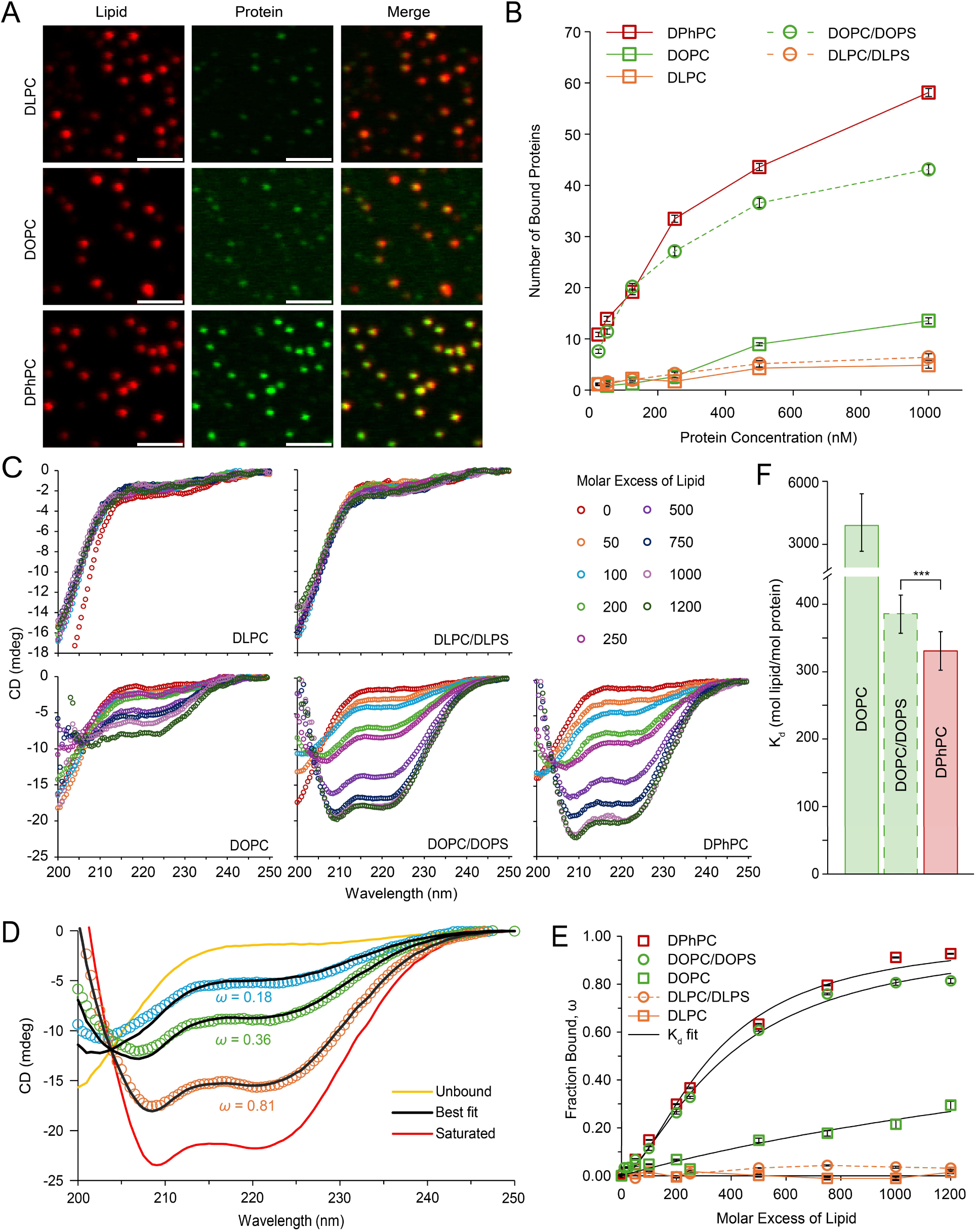
Packing defects are necessary and sufficient for NTA-αSyn binding regardless of membrane charge. (A) Lipid, protein, and merged fluorescent images of SUVs incubated with 500 nM NTA-αSyn. Scale bars represent 2 µm. (B) Binding curves for NTA-αSyn to SUVs with an average diameter of 100 nm. This data was obtained from fluorescence microscopy. N = 900-3554 vesicles for each point on the binding curve. Error bars represent the 99% confidence interval of the mean. (C) CD spectra of NTA-αSyn for each lipid composition. Molar excess of lipid is the lipid molar concentration divided by protein molar concentration. A protein of 5 µM was used for these experiments. (D) Representative plots illustrating the extrema CD spectra used for fitting and determining intermediate ω values. For these representative spectra, sonicated 3:1 DPhPC:DPhPS vesicles were used. (E) Binding curves for unlabeled NTA-αSyn obtained from CD spectroscopy. Error bars represent the standard error of the mean (SEM) for the fitting parameter, ω. (F) Dissociation constant (Kd) obtained from fitting the data in panel D to Equation 2. All PC/PS membranes contained a 3:1 PC:PS molar ratio. *** corresponds to P < 0.001 as determined by Student’s t-test.

These trends were mirrored in binding curves that were obtained using CD spectroscopy (**Figure 3C**). Here, we directly monitored NTA-αSyn insertion into membrane defects by tracking spectral changes of the non-fluorescently labeled protein as it transitioned from a completely disordered structure to a partially folded a-helical structure upon membrane binding ^48,55^. To limit curvature-influenced effects, SUVs were prepared such that the vesicle size distributions were comparable among all vesicle compositions (**Figure S6**). After CD spectra were obtained for lipid- free αSyn and lipid-saturated αSyn, intermediate spectra were fit using **Equation 1** to determine ω, the fraction of αSyn bound to vesicles for a given lipid concentration (**Figure 3D, Figure S6, and Table S3**). Here, the lipid concentration is reported as a molar excess of lipid (i.e., moles of lipid per mole of protein). In **Figure 3E** ω is plotted as a function of the lipid concentration or 5 different compositions. The mirrored binding curves in **Figures 3B and 3E** verify that our fluorescence microscopy-based tethered vesicle assay was appropriate for measuring membrane insertion of αSyn. The binding curves in **Figure 3E** were fit to a Langmuir-Hill binding curve (**Equation 2**) to determine an apparent dissociation constant (Kd) for each composition (**Table S4**). Kd could not be determined for DLPC and DLPC/DLPS systems due to the exceptionally low binding observed. Of the DOPC, DOPC/DOPS, and DPhPC systems analyzed in **Figure 3F** Kd was lowest for DPhPC membranes and highest for DOPC membranes, further indicating that binding affinity was highest for DPhPC membranes.

### Negative surface charge synergistically enhances NTA-**α**Syn binding to membranes

We next investigated whether negative charge affects NTA-αSyn binding to DPh membranes, which are inherently defect-rich. To carry out these experiments, we utilized diphytanoylphosphatidylserine (DPhPS) mixed with DPhPC at a molar ratio of 3:1 DPhPC:DPhPS (**Figure 4**). The tethered vesicle assay (**Figures 4A and S4**) and CD spectroscopy (**Figures 4B and S7**) both revealed that NTA-αSyn exhibited preferential binding to membranes containing 3:1 DPhPC:DPhPS in comparison to membranes containing just DPhPC. Of note, the inclusion of PS increased the surface coverage of packing defects in DPh membranes by increasing the average size of individual membrane defects (**Figure 4C**). When comparing the results of PC experiments and PC/PS experiments as a function of lipid tail chemistry, we found that the binding of NTA-αSyn exhibited a strong linear response to surface coverage by defects with both fits intersecting the x-axis at a value of approximately 2.5% (**Figure 4D**). This strong linear response is indicative of binding in the sub-saturation regime, where the amount of binding is directly proportional to the number of binding sites available.

**Figure 4.**
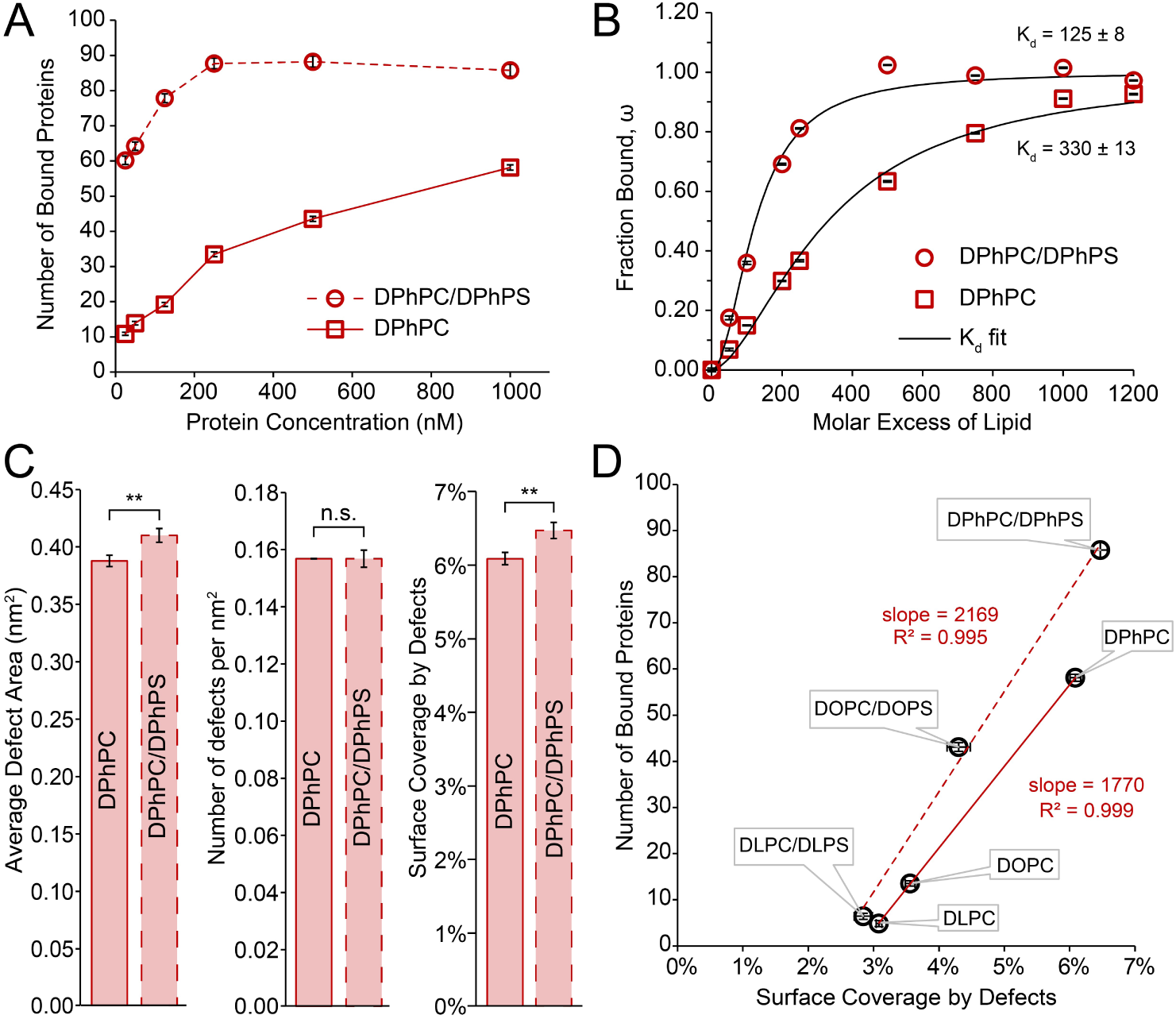
Packing defects and anionic charge synergistically enhance NTA-αSyn binding to membranes. (A) Binding curves for NTA-αSyn obtained from microscopy on vesicles with an average diameter of 100 nm. Each point represents anywhere between 1307 – 3554 analyzed vesicles. Error bars represent the 99% confidence interval of the mean. (B) Binding curves for NTA-αSyn obtained from CD spectroscopy. Error bars represent the SEM for the regression parameter ω at each protein:lipid ratio. (C) Average defect area (left), defect frequency (center), and percentage coverage of the membrane surface by packing defects (right) as determined by molecular dynamics simulations. Error bars represent the SD (N=3). (D) The measured binding response of NTA-αSyn to packing defects for 100% PC membranes and 3:1 PC:PS membranes. These binding responses correspond to 100- nm diameter SUVs that were incubated with 1 µM NTA-αSyn in tethered vesicle experiments. For DPhPC and DPhPC/DPhPS, the horizontal and vertical error bars are taken from C and A, respectively. The horizontal and vertical error bars for all other compositions are taken from Figure 2E and Figure 3B, respectively. ** corresponds to P < 0.01 as determined by Student’s t-test.

### NTA-**α**Syn binding can be modulated by the degree of lipid unsaturation

We next sought to determine whether the density of membrane packing defects could be modulated by lipid tails that are more relevant than DPh in physiological contexts. Due to αSyn’s localization in the brain, the compositional intricacies of both neuronal membranes and synaptic vesicles must be considered. PUFAs, for example, are enriched in both neurons ^56^ and synaptic vesicles ^53,57^. Since PUFA lipid tails occupy more space than saturated lipid tails for a given phospholipid, we hypothesized that PUFA tails could be used as a means of changing the density of packing defects and subsequently, the amount of NTA-αSyn binding. To test the impact of lipid unsaturation on NTA-αSyn binding, we synthesized and monitored binding to vesicles containing 40/25/25/10 mol% cholesterol/PC/PE/PS to mimic the naturally occurring lipid composition in synaptic vesicles ^53^ (**Figure 5**). Here, PE refers to lipids containing phosphatidylethanolamine headgroups. Lipid tails for PE and PS lipids were composed of oleic acid (i.e., DOPE and DOPS). Lipid tails within the PC component were any of the following: oleic acid (i.e., DOPC, also known as 18:1 PC), linoleic acid (18:2 PC), arachidonic acid (20:4 PC), or docosahexaenoic acid (22:6 PC) (**Figure 5A**). Our simulation results revealed that the surface coverage by membrane defects indeed increased as the degree of lipid unsaturation increased (**Figure 5B**). Interestingly, there was a concomitant increase in NTA-αSyn binding with an increase in the degree of unsaturation as revealed using fluorescence microscopy (**Figure 5C**). These results were also observed when SUVs were incubated with lower concentrations of NTA-αSyn (**Figure S8**). Similar to the results in **Figure 4D**, this binding exhibited a direct proportionality to surface coverage by membrane defects but notably intersects the x-axis at a value near 0% (**Figure 5D**).

**Figure 5.**
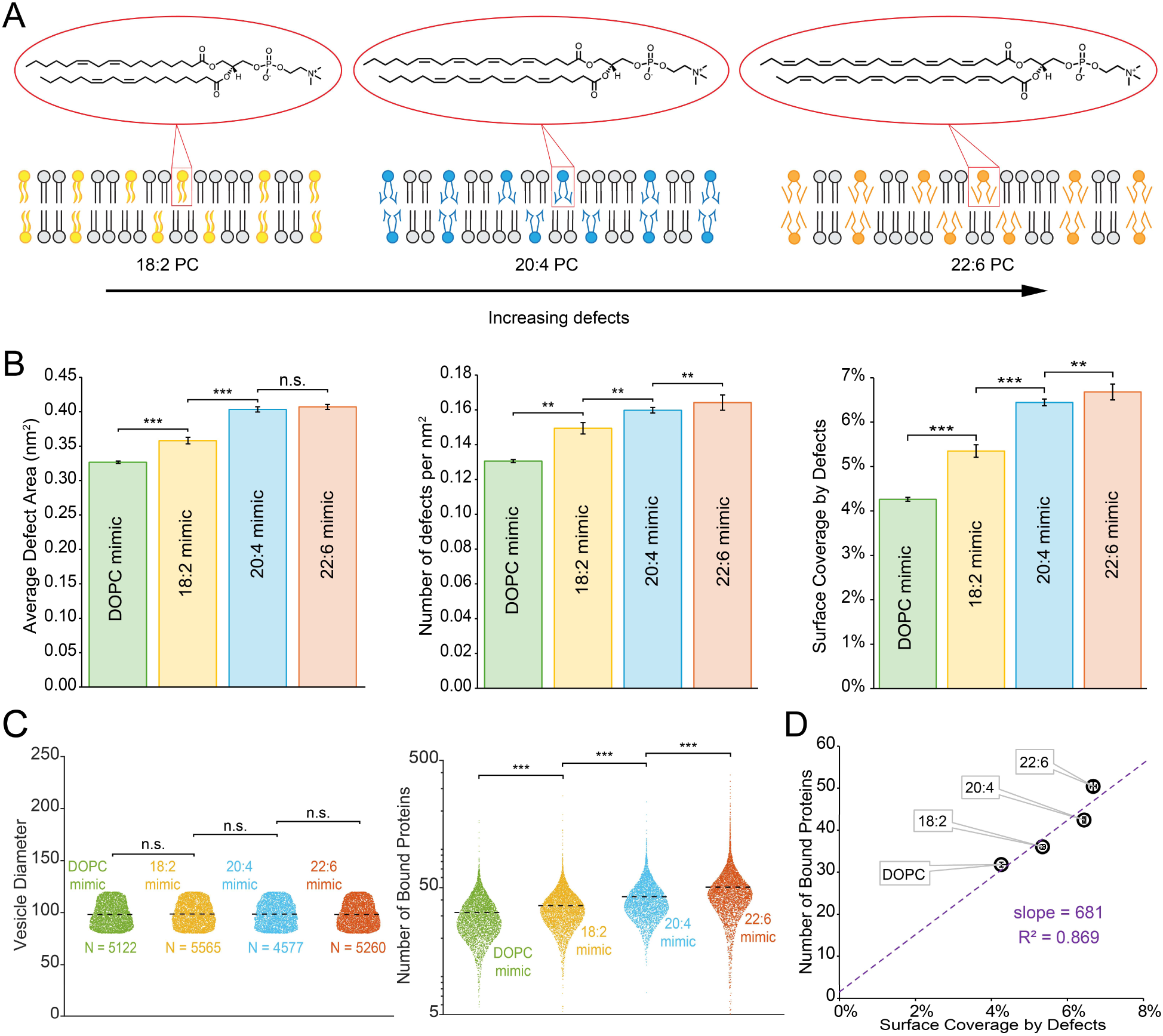
Lipid polyunsaturation modulates membrane packing defects in synaptic vesicle mimics, which in turn regulates NTA-αSyn binding. (A) Chemical structures of the PUFAs used in the synaptic vesicle mimics and schematics of lipid bilayers containing these PUFAs. As the degree of unsaturation increases, the tails become increasingly bulkier and exhibit increasing steric repulsion that leads to more packing defects. (B) Average defect area (left), defect frequency (center), and percentage coverage of the membrane surface by packing defects (right) as determined by molecular dynamics simulations. Error bars represent the SD (N=3). (C) The distribution of diameters for analyzed vesicles that were incubated with 1 µM αSyn (left). Number of proteins bound to vesicles incubated with 1 µM NTA-αSyn (right). The black dashed line indicates the average values of the data sets. For each composition, a small number of vesicles were omitted from these plots as they fell below the minimum y-axis value of 5 proteins bound. The numbers of vesicles omitted are as follows: DOPC mimic (4 out of 5122 vesicles), 18:2 mimic (8 out of 5565 vesicles), 20:4 mimic (1 out of 4577 vesicles), and 22:6 mimic (29 out of 5260 vesicles). (D) The measured binding response of NTA-αSyn to packing defects for synaptic vesicle mimics. These binding responses correspond to the data from panels B-C. The horizontal error bars are taken from B, and the vertical error bars represent the 99% confidence interval of the mean from each composition in panel C (right). For panel B, ** and *** correspond to P < 0.01 and P < 0.001, respectively, as determined by Student’s t-test. For panel C, *** corresponds to P < 0.0001 as determined by unpaired Student’s t-test.

## Discussion

αSyn’s relevance to PD and other neurodegenerative disorders has been well-studied, and the membrane binding of αSyn has been previously shown to be an essential part of its pathology through the nucleation of aggregation and fibrillization ^9,13,14^. Here, we provide insights into αSyn’s binding to reconstituted membrane surfaces by modulating the lipid compositions, revealing that membrane packing defects are the primary driver of αSyn binding. Our results demonstrate that NTA-αSyn, the physiologically prevalent form of αSyn, has greater binding to defect-rich, neutrally charged surfaces composed of DPhPC over anionic membranes with fewer packing defects, such as those containing DOPC:DOPS at a 3:1 molar ratio. NTA may selectively enhance αSyn’s binding to DPhPC membranes by increasing its inherent α-helical content in the unbound state ^49,58^, thus reducing the energy barrier necessary for membrane insertion ^50^. NTA also increases the hydrophobicity of the N-terminus of αSyn, replacing the charged amine with a neutral acetyl group, incentivizing additional hydrophobic interactions ^59^. While this reduction in positive charge diminishes electrostatic interactions between the unmodified, N-terminal methionine and anionic lipid headgroups, recent MD simulations have shown that this loss in electrostatic contribution to binding energy is offset by increased hydrophobic contributions facilitated by the acetylated methionine ^58^. These simulations also demonstrate that acetylation promotes extension of αSyn’s helical structure along the length of the N-terminus, enhancing membrane binding. This state of increased binding activity would, therefore, be highly amenable to defect-rich membranes such as DPhPC, as these membranes contain higher exposure of the underlying lipid tails. It is relevant here to note that we do not evaluate the specific alpha-helical conformation (i.e. broken helix vs extended helix) of αSyn’s N terminus upon membrane binding in this work. This precise structural conformation of the helical N terminus in the membrane-bound state remains context-dependent and lacks strong consensus, as it depends on membrane size and curvature ^60,61^. In **Figure 1E**, αSyn is depicted as an extended helix, which is the likely conformation adopted upon binding to 100 nm vesicles ^62^.

The observed preference for membrane defects by NTA-αSyn was preserved across both binding assays used. Fluorescence microscopy and circular dichroism spectroscopy combined with all-atom molecular dynamics simulations demonstrated that the presence of defects dictates the ability of αSyn to insert into membranes, and that negatively charged membranes cannot overcome a lack of membrane defects necessary for αSyn binding. This result is particularly interesting in the context of recent MD simulations that have probed the relative contribution of electrostatic and hydrophobic forces for αSyn membrane binding ^58,63–66^. This literature jointly describes a multi-state binding event in which electrostatic and hydrophobic interactions serve distinct roles in αSyn binding. Electrostatics dominate when the negative headgroups interact with the lysine-rich N-terminus of αSyn, bringing the protein close to the membrane surface. The hydrophobic interactions then dominate when the N-terminus embeds into the membrane upon binding. These hydrophobic interactions become even more relevant in the context of NTA-αSyn where MD simulations show additional helical propensity increases membrane binding affinity ^58^. Throughout this literature, electrostatic interactions were shown to assist in αSyn binding through membrane recognition, but the defects and hydrophobic interactions mediate the physical insertion of the protein beyond surface adsorption ^66^. This necessity of defects for binding and insertion was exemplified with our results utilizing DL membranes, where the addition of 25% charge from PS did not increase the density of packing defects nor the amount of bound αSyn. Even in the presence of substantial anionic charge, which assists with αSyn localization to the membrane, insufficient defects resulted in negligible binding to DL membranes. In contrast, we note a substantial increase in defect coverage and binding of αSyn to DO membranes when 25% PS was included. This increase in defect content could be due to repulsion between charged lipid headgroups that open additional defects in the DO system or due to a mismatch in headgroup size between PC and PS ^59^. While increased αSyn binding was strongly correlated with an increase in defect content and neutral DPhPC membranes elicited stronger binding than anionic DOPC/DOPS membranes, it must be noted that anionic charge and defect content exhibited synergy in recruitment of αSyn. Interestingly, our CD Spectroscopy results suggest that the addition of PS enhanced cooperativity of αSyn binding in both DO and DPh membranes (**Table S4)**.

Notably, αSyn exhibited strong linear binding responses on pure PC membranes, PC/PS membranes, and cholesterol-containing synaptic vesicle mimic membranes. Interestingly, PC and PC/PS membranes yield similar x-intercept values of approximately 2.5%, indicating that there is a minimum coverage of defects necessary for αSyn to bind to these membranes. This explains why, when PS lipids were added to the DL system, there was a lack of binding response from NTA-αSyn as DL defect coverages were near this minimum value of approximately 2.5% (**Figure 2C**). When comparing the neutral PC systems to the anionic PC/PS systems in **Figure 4D**, the anionic membrane systems had a steeper slope than the neutral membrane systems. This steeper slope indicates that for a given increase in defect content, anionic membranes elicit a greater increase in NTA-αSyn binding. This result correlates well with existing literature stating the importance of anionic charge for αSyn binding and further emphasizes synergistic behavior between packing defects (i.e., hydrophobic interactions) and membrane charge (i.e., electrostatic interactions) for αSyn binding. In contrast to the defect minimums seen in the PC membranes (0% PS) and PC/PS membranes (25%), the linear trend for synaptic vesicle mimicking membranes in **Figure 5D** exhibits a much lower slope with an x-intercept of approximately 0%. This result was somewhat surprising considering that these membranes contained 10% PS, which would be expected to yield an intermediate slope to the 0% PS and 25% PS systems. This lower slope could be attributed to either the inclusion of PE lipids or cholesterol. It is known that adding DOPE to DOPC/DOPS mixtures increases αSyn binding in SUVs^27^, so it is likely that the addition of cholesterol is reducing NTA-αSyn’s response to increasing packing defects as adding cholesterol tightens membranes, reducing fluidity, and reducing overall αSyn membrane affinity ^18^. Other studies have shown cholesterol reduces αSyn’s affinity to membranes by either reducing the interaction of the second half of the N-terminal α-helix with membranes ^67^ or attenuating the electrostatic interaction between the membrane and αSyn ^66^. In the context of our results, some combination of these mechanisms could be reducing αSyn binding at higher defect content, but it is likely that membrane fluidity reduction and tightened packing is a prominent mechanism with 40 mol% cholesterol membranes. Interestingly, our extrapolation in **Figure 5D** suggests that cholesterol-containing membranes promote stronger NTA-αSyn binding in the low defect regime than the pure PC and PC/PS membranes used in **Figure 4D**. NTA-αSyn binding to cholesterol-rich compositions with low defects could be due to cholesterol inclusion loosening the membrane in densely packed compositions allowing for αSyn insertion ^64^ or the direct interaction between the hydroxyl headgroup and αSyn ^18^. Additionally, recent research has shown that αSyn can serve as a cholesteryl-ester (CE) specific sensor in lipid droplets and bind specifically to a pure CE lipid droplet ^68^. This type of preferential cholesterol interaction may contribute to membrane recruitment of NTA-αSyn in the low defect regime. Taken together, it seems that cholesterol serves as a complementary regulator of αSyn binding. While membrane defects are necessary for αSyn helix insertion, cholesterol could strengthen αSyn binding in defect-poor compositions through a cholesterol specific interaction and attenuate αSyn binding in defect-rich compositions.

The inclusion of PUFAs into synaptic vesicle mimics provides insight into how cells might regulate αSyn binding in physiological contexts. We observed a strong correlation between αSyn binding to membranes and the degree of unsaturation in lipid tails. Incidentally, the enrichment of PUFAs in the brain leads to the formation of oligomeric αSyn ^51,52,69,70^. More specifically, the 22:6 PUFA, which is the PUFA that elicited the highest degree of αSyn binding in our synaptic vesicle mimic experiments, is known to be further enriched in PD patients with a distinct link in enhancing αSyn neuropathology and oligomerization ^25,71–74^. We, therefore, hypothesize that this enrichment of PUFAs in the brain may lead to the formation of oligomeric αSyn by promoting higher seeding densities for αSyn on the membrane surface. This proposed mechanism indicates how lipid polyunsaturation could regulate membrane packing defects, thus impacting αSyn interactions with synaptic vesicles and, by extension, aSyn pathology.

Previous studies have neglected the significance of αSyn interactions with zwitterionic membranes, citing evidence of reduced binding compared to anionic membranes. However, this current study, along with a growing body of work ^11,12,32^, shows that αSyn interactions with zwitterionic membranes, when sufficient defects are present, can be comparable if not greater than a charged membrane. Following our results, future studies should reframe analyses that focus solely on anionic membranes to include interactions with zwitterionic membranes. For example, αSyn’s direct interactions with the endoplasmic reticulum (ER), a noted zwitterionic membrane^75^, have been shown to impact vesicle trafficking and increase ER stress leading to neuroinflammation and neurodegeneration^76^. Although these types of ER interactions have been observed, the underlying cause remains unexplained in the literature. Our study fills this knowledge gap by elucidating mechanisms by which physiological αSyn interacts with electrostatically neutral membranes. Additionally, our work informs molecular mechanisms for therapeutic intervention. Previous work has shown that αSyn fibrillization and toxicity can be reduced in dopaminergic neurons by incubating them with zwitterionic lipid nanoparticles (LNPs) ^77^. While the authors considered their findings controversial in the context of αSyn-membrane literature, they hypothesized that αSyn interacts with zwitterionic LNPs to reduce the cytosolic pool of free αSyn available for fibrillation and aggregation. Our current study corroborates these findings by experimentally demonstrating that αSyn can preferentially bind to zwitterionic membranes when the tail chemistry of the phospholipids is favorable. Future work in therapeutic applications could leverage our findings to design and formulate LNPs with enhanced specificity for αSyn interactions.

## Conclusion

In conclusion, we show that membrane packing defects are the primary drivers of NTA-αSyn binding in both neutral and anionic phospholipid membranes. Interestingly, NTA-αSyn did not exhibit appreciable binding to anionic membranes until a critical threshold of packing defects was reached. While anionic charge did synergistically enhance NTA-αSyn binding above this minimum defect threshold, these enhancements were coupled to a further increase in membrane defect content likely resulting from electrostatic repulsion between phospholipid head groups. Furthermore, we saw that αSyn binding can be modulated by the degree of unsaturation in lipid tails, which directly controls the density of membrane packing defects. We also noted that cholesterol differentially regulates the membrane binding of NTA-αSyn, enhancing binding in low-defect regimes while inhibiting it in the high-defect regimes. Our results outline molecular mechanisms by which lipid compositions can regulate membrane-nucleated oligomerization and pathology of αSyn. These findings necessitate a reevaluation of the current framework surrounding αSyn-membrane interactions as previous studies primarily focused on anionic membranes. By challenging the prevailing notion that electrostatic interactions with lipid headgroups primarily drive αSyn-membrane interactions, this work underscores the necessity of considering lipid tail chemistry when investigating αSyn- membrane interactions. This mechanism of binding through membrane packing defects likely extends beyond αSyn, pointing to a broader paradigm in protein-membrane interactions. By emphasizing lipid tail chemistry and membrane defect density, this study may uncover how various proteins detect and adapt to specific membrane environments, deepening our understanding of membrane-driven processes.

## Materials and Methods

### Materials

1,2-dioleoyl-sn-glycero-3-phosphocholine (DOPC), 1,2-dioleoyl-sn-glycero-3-phospho-L-serine (sodium salt) (DOPS), 1,2-dioleoyl-sn-glycero-3-phosphoethanolamine (DOPE), 1,2-diphytanoyl-sn-glycero-3-phosphocholine (DPhPC), 1,2-diphytanoyl-sn-glycero-3-phospho-L-serine (sodium salt) (DPhPS), cholesterol (plant), 1,2-dilauroyl-sn-glycero-3-phosphocholine (DLPC), 1,2-dilauroyl-sn-glycero-3-phospho-L-serine (sodium salt) (DLPS), 1,2- dilinoleoyl-sn-glycero-3-phosphocholine (18:2 PC), 1,2-diarachidonoyl-sn-glycero-3-phosphocholine (20:4 PC), and 1,2-didocosahexaenoyl-sn-glycero-3-phosphocholine (22:6 PC) were purchased from Avanti Polar Lipids. Dipalmitoyl-decaethylene glycol-biotin (DP-EG15-biotin) was generously provided by D. Sasaki from Sandia National Laboratories, Livermore, CA ^78^. ATTO 647N 1,2-Dipalmitoyl-sn-glycero-3-phosphoethanolamine (DPPE) and ATTO 488-maleimide were purchased from ATTO-TEC. Biotin-PEG-Silane (5000 MW) and mPEG-Silane (5000 MW) were purchased from Laysan Bio, Inc. Isopropyl-beta-D-thiogalactoside (IPTG) was purchased from Gold Biotechnology. Tris(2-carboxyethyl) phosphine hydrochloride (TCEP), phenylmethanesulfonyl fluoride (PMSF), sodium phosphate monobasic, sodium phosphate dibasic, and β-mercaptoethanol (BME) were purchased from Sigma-Aldrich. 4-(2-hydroxyethyl)-1-piparazineethanesulphonic acid (HEPES) and tris(hydroxymethyl)aminomethane (Tris) were purchased from Fisher Scientific.

### **α**-Synuclein Expression and Purification

The wild-type α-synuclein expression plasmid (pET21a backbone) was provided by Peter Chung (University of Southern California, Los Angeles, CA). The mutated (Y136C) α-synuclein expression plasmid (pRK172 backbone) was provided by Jennifer C. Lee (National Institutes of Health, Bethesda, MD). Each respective plasmid was transformed into BL21 (DE3) cells that were pre-transformed with the pNatB plasmid coding for *N*-α- acetyltransferase to acetylate the α-synuclein ^79^. After incubation in 2xTY media at 37° C to OD600 ∼ 0.6, cells were induced with 1mM IPTG and incubated for an additional 4 hours. Cells were pelleted and resuspended in 25 mM Tris, 20 mM NaCl, 1 mM PMSF, and 1x Roche protease inhibitor cocktail (Sigma-Aldrich) (pH 8.0). 10 mM BME was added for the Y136C mutant to prevent disulfide bonding. The cells were sonicated on ice at 65% amplitude for 15 minutes, and then 30uL/10mL of cell solution of DNAse I (Thermo Fisher Scientific) was added to the solution and incubated for 30 minutes at 37° C. The pH of the solution was reduced to pH 3.5, and the cell debris was pelleted for 25 minutes at 20,000 xg. The pH of the supernatant was set to pH 7.0, and a 50% ammonium sulfate (Sigma Aldrich) precipitation was performed with the protein precipitating out and pelleting after another 25 minute, 20,000 xg centrifugation. The pellet was resuspended in 20mM Tris (pH 8.0), run over a 5mL HiTrap Q FF column (Cytiva), and eluted with 20mM Tris, 1M NaCl (pH 8.0) with 10mM BME in both buffers for the Y136C mutant. The eluted protein was concentrated using a 3K MWCO centrifugal filter (Sigma-Aldrich) and run through SEC on a Superdex 200 10/300 column (Cytiva) into 25mM HEPES, 150mM NaCl (pH 7.4) with 0.5mM TCEP for the Y136C mutant. The appropriate fractions were concentrated, and the final concentration was measured using UV-Vis spectroscopy at 280 nm wavelength. The extinction coefficients used were 5960 M^-1^ cm^-1^ and 4470 M^-1^ cm^-1^ for WT-αSyn and Y136C-αSyn, respectively.

### **α**-Synuclein Fluorescent Labeling

ATTO 488-maleimide was stored as 100 mM aliquots in DMSO at -80°C. Labeling was performed by combining ATTO-488 maleimide with Y136C-αSyn at a 6x molar excess of dye. The reaction was allowed to proceed overnight at 4° C in the dark. The total amount of DMSO added to Y136C-αSyn never exceeded 2% v/v. Excess dye was removed by flow through a Superdex 75 10/300 GL (Cytiva) size exclusion chromatography column. The protein:dye labeling ratio was confirmed to be 1:1 by UV-Vis spectroscopy. An extinction coefficient of 90000 M^-1^ cm^-1^ was used for ATTO 488 at 488 nm, with a correction factor of 0.09 for the 280 nm / 488 nm absorption ratio.

### SUV Preparation

Lipid aliquots were thawed from -80°C and combined in the following molar percentages for use in the tethered vesicle assays:

- 97.5% DPhPC, 2% DP-EG15-biotin, 0.5% ATTO 647N
- 72.5% DPhPC, 25% DPhPS, 2% DP-EG15-biotin, 0.5% ATTO 647N
- 97.5% DOPC, 2% DP-EG15-biotin, 0.5% ATTO 647N
- 72.5% DOPC, 25% DOPS, 2% DP-EG15-biotin, 0.5% ATTO 647N
- 97.5% DLPC, 2% DP-EG15-biotin, 0.5% ATTO 647N
- 72.5% DLPC, 25% DLPS, 2% DP-EG15-biotin, 0.5% ATTO 647N
- 23.75% DOPC, 23.75% DOPE, 10% DOPS, 40% Cholesterol, 2% DP-EG15-biotin, 0.5% ATTO 647N
- 23.75% 18:2 PC, 23.75% DOPE, 10% DOPS, 40% Cholesterol, 2% DP-EG15-biotin, 0.5% ATTO 647N
- 23.75% 20:4 PC, 23.75% DOPE, 10% DOPS, 40% Cholesterol, 2% DP-EG15-biotin, 0.5% ATTO 647N
- 23.75% 22:6 PC, 23.75% DOPE, 10% DOPS, 40% Cholesterol, 2% DP-EG15-biotin, 0.5% ATTO 647N

The following molar percentages were used for vesicle preparation in CD spectroscopy experiments:

- 100% DOPC
- 75% DOPC, 25% DOPS
- 100% DPhPC
- 75% DPhPC, 25% DPhPS
- 100% DLPC
- 75% DLPC, 25% DLPS

Solvents were evaporated with a nitrogen stream, followed by vacuum storage for at least 2 hours. Dried lipid films were rehydrated using 20mM sodium phosphate and 150mM NaCl (pH 7.0) buffer. After 5 minutes of hydration, the lipid suspensions were subjected to 3 freeze/thaw cycles by submerging the solution in liquid nitrogen for 2 minutes and thawing at 60° C for 3 minutes. The SUVs were then extruded through a 50nm polycarbonate membrane (Whatman). The average diameter of vesicles used for CD spec across all compositions was 67 ± 5 nm, determined by dynamic light scattering (**Fig. S6**). When handling PUFA-containing mixtures, special precautions were taken to minimize oxidation. All PUFA aliquots were handled in a N2 glove box, and all buffers were purged with N2 before rehydrating the lipid films. Additionally, freeze-thaw cycles were omitted to limit exposure to any residual oxygen, and lipid extrusion was carried out within a N2 glove box. Samples were then imaged within 30 minutes of air exposure. All other vesicle preparations were imaged within 48 hours of preparation and stored in a 4 °C refrigerator to minimize degradation of quality. Unilamellar structures were verified with a volume labeling assay (Figure S9), and their surface charge was confirmed with a PLL adsorption assay (Figure S10).

### Slide Passivation and Tethered Vesicle Assay

Glass coverslips were cleaned with a modified RCA protocol and passivated with PEG-silane, as described previously, with slight modifications ^80^. Briefly, glass slides were soaked in a concentrated KOH/peroxide bath, followed by a concentrated HCl/peroxide bath each at 80 °C for 10 minutes. Once cleaned, the slides were passivated by surface treatment with a 7.5 mg/mL PEG-silane solution in isopropanol. The solution was comprised of 5% biotin-PEG-silane (5000 MW) and 95% of mPEG-silane (5000 MW). All materials for creating the PEG-silane solution were handled in an N2 environment. Immediately before passivating the slide, 1% v/v of glacial acetic acid was added to catalyze the reaction between silanes and hydroxyl groups on the cover slip. 55uL of the solution was added to the top surface of the cleaned coverslips, spread out, then placed into an oven at ∼ 70 °C for 30 - 60 minutes. The slides were then rinsed under DI water to remove excess PEG-silane and dried under N2. These slides were stored dry under air for a maximum of 1 week.

Imaging wells were created by placing 0.8 mm thick silicone sheets (Grace Bio-Labs) with 5mm holes on the passivated slide. Each well was hydrated with 6 μg of NeutrAvidin (ThermoFisher) in 30μL of 20mM sodium phosphate, 150mM NaCl (pH 7.0) and incubated for 10 minutes. After incubation, each well was rinsed to remove excess NeutrAvidin. Following this step, each well was rinsed with a solution containing SUVs to reach a 1μM lipid concentration. After 10 minutes of incubation, excess SUVs were rinsed from the wells with the same buffer, and the α-synuclein tagged with ATTO 488 was added at the appropriate concentration.

### Confocal Microscopy

Imaging was performed on a laser scanning confocal microscope (Leica Stellaris 5). Two excitation lasers were used: 488 nm and 638 nm for labeled αSyn and SUVs, respectively. The wavelength detection bandwidths used were 493 – 558 nm and 643 – 749 nm for labeled αSyn and SUVs, respectively. A Leica HC PL APO 63x, 1.4 NA oil-immersion objective was used to acquire images, and the zoom factor was set such that square pixel sizes of 70 nm were obtained. All images were acquired using a scan speed of 400 Hz.

### Image Processing for the Tethered Vesicle Assay

Image processing was performed similarly to previous work ^81^. Briefly, images of diffraction-limited, fluorescent lipid and protein puncta were acquired using confocal microscopy. Each image was a three-frame repeat of a single field of view. Using publicly available particle detection software (cmeAnalysis), the 2D fluorescence distribution of all puncta in each image was fit to 2D-gaussian functions ^82^. Using this fit, the fluorescent amplitudes were determined for the puncta in both the lipid and protein channels. Only puncta in the lipid channel that persisted through all three consecutive image frames in a single field of view were considered in this analysis, ensuring that all vesicles detected were above the fluorescent noise threshold. As an additional filtering step, any puncta whose amplitude was not significantly above the level of background fluorescence was excluded from the analysis. Our analysis also excluded vesicles with diameters above 250 nm to ensure that all analyzed puncta were truly diffraction-limited. Protein binding for each vesicle was determined by analyzing the colocalization between both fluorescent channels. The search radius for colocalized protein fluorescence amplitudes was three times the standard deviation of the Gaussian fit of the point spread function in the lipid channel.

### Calibration of Vesicle Diameters and Number of Bound Proteins

SUV diameters and the number of proteins bound to each vesicle were determined as previously described ^51,81^. Briefly, tethered SUVs were imaged to obtain a distribution of the fluorescence intensities for the vesicle population. A conversion factor was then determined by overlaying the square root of the fluorescence distribution and the DLS diameter distribution (**Fig. 1B**). All tethered vesicles included in the binding analysis (**Fig. 1**, **Fig. 3-5, Fig. S2-S4, S8-S9**) were subsets of their respective vesicle populations and selected in the diameter range 80-120 nm, resulting in average diameters of 100 nm across all compositions studied. This step was necessary to control for curvature-sensing effects in the interpretation of our results.

To determine the number of proteins bound to each vesicle, we performed single-molecule imaging of fluorescent αSyn that was adsorbed to glass coverslips. The fluorescence intensities of the punctate structures were used to generate a distribution (**Fig. 1B**). The peak value of this fluorescence intensity distribution was then used to represent the fluorescence intensity of a single ATTO 488 labeled αSyn. Quantitative adjustments were made during the data processing to account for changes in image frame accumulations and linear gain settings on the detector.

### Circular Dichroism Spectroscopy

Circular dichroism measurements were collected on a JASCO J-1100 instrument with a 1 mm path length quartz cuvette (JASCO). Scans were taken from 195 – 300nm at a 0.5 nm data pitch with 100 nm/min scan speed. The data integration time was 2 seconds, and each scan was accumulated 2 times. The buffer used for each experiment was 20 mM sodium phosphate, 150mM NaCl (pH 7.0). Samples of vesicles and protein were allowed to equilibrate for at least 15 minutes before spectra were acquired. Each sample contained 5 μM αSyn and was blanked by a buffer solution containing SUVs at the appropriate lipid concentration.

Equation 1 was used to determine ω, the fraction of bound αSyn. Here, CU is the average spectrum for unbound, disordered αSyn, CS is the average saturated spectrum for membrane-bound αSyn, and CI is the intermediate spectrum for αSyn that is partially bound to membranes (i.e., a mixture of bound and unbound states). Sonicated SUVs composed of 3:1 DPhPC/DPhPS were used to obtain saturated spectra, CS.

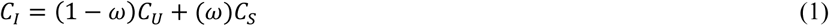

The apparent dissociation constant, Kd, was found using a Langmuir-Hill adsorption isotherm to fit ω as a function of the lipid:protein molar ratio, [L:P] (Equation 2). Here, n represents the Hill coefficient.

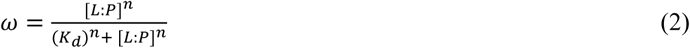

### All-Atom Molecular Dynamics Simulations

Experimental and mathematical details for all simulations performed are described in detail in the Supplementary Information.

## Acknowledgments

Thank you to Jennifer C. Lee for generously providing the pRK172 human αSyn(Y136C) plasmid. This research was supported by the National Institutes of Health through R35GM147333 to D.H.J., O.H.K., and W.F.Z, and R35GM150716 to A.M. and P.J.C.

