## Supplementary Information for "Lipid Packing Defects are Necessary and Sufficient for Membrane Binding of α-Synuclein"

### Simulation Methods

**Atomistic Models.** Atomistic lipid membrane models were built using the CHARMM-GUI [1,2] webserver's Membrane Builder [3] functionality. System geometry, composition, and simulation parameters were inspired by Garten and coworkers who performed similar analysis on other lipid membrane simulations [4]. **Table S1** lists the composition of each model system. The CHARMM36 force field was used to model lipid and ion interactions [5,6]. CHARMM TIP3P was used to model water interactions. Cholesterol and all lipids other than DPhPS were pre-defined within the CHARMM-GUI database of lipids and were used without modification. A force field for DPhPS was developed within the CHARMM-GUI ligand modeler [7] using the CHARMM General Force Field [8] (CGenFF v 4.6). Because DPhPS is not native to CHARMM-GUI's Membrane Builder, the initial configuration of DPhPS-containing systems was alternatively generated using Packmol [9]. In both Packmol and Membrane Builder, 50 water molecules per lipid molecule were added along with $Na^+$ and $Cl^-$ ions to bring the salt concentration to 150 mM. The number of water molecules was chosen to match the protocol of Garten and coworkers [4] who previously analyzed structures of lipid membranes and their defects. Standard treatment of electrostatics in MD simulations requires the system to be net neutral in charge. Therefore, additional $Na^+$ or $Cl^-$ ions were used to neutralize the box in systems where charged lipids were used. **Figure 2** illustrates two of the model systems. While the vesicles used in this work had average diameters of ~100 nm and the simulations were performed on flat bilayers, previous work has verified that defect size increases by only ~10% on 100 nm vesicles when compared to flat bilayers, regardless of lipid composition [10]. Therefore, it can be assumed that the flat bilayers simulated in this work serve as accurate representations of their vesicular counterparts with 100 nm diameters.

**MD Simulation Parameters.** All MD simulations presented here were conducted in GROMACS 2021.6 [11]. A standard Lennard-Jones 12-6 model with Lorentz-Berthelot mixing rules was used to for non-bonded interactions. Lennard-Jones interaction potentials were cut off at a distance of 1.2 nm with a force-switch modifier applied from 1.0 to 1.2 nm. Electrostatic interactions were calculated explicitly in normal space at short distances. Long-range electrostatic interactions were calculated using the particle mesh Ewald method. By default, GROMACS automatically adjusts the short-range electrostatic cutoff distance to optimize performance while maintaining a low error. The Verlet leap-frog integrator was used to integrate the equations of motion. An integration timestep of up to 2 fs was facilitated by constraining bonds between hydrogen atoms and heavy atoms using the LINCS algorithm. Thermostats and barostats were used to control temperature and pressure. During all steps, a velocity rescaling (v-rescale) thermostat was used. During some steps of equilibration, a cell-rescaling (c-rescale) barostat was used. A Parinello Rahman barostat was used for production simulations. Temperature was maintained at its set point of 303.15K with a time constant of 1 ps. Regardless of which barostat was used, pressure was maintained at 1 bar with semi-isotropic coupling and a time constant of 5 ps and a compressibility of $4.5 \times 10^{-5}$ $bar^{-1}$. Net motion of the center of mass of the system was removed every 200 fs.

**Minimization and Equilibration.** In addition to generating starting configurations, CHARMM-GUI provides a set of suggested parameters for energy minimization and six steps of equilibration. The suggested parameters were used as-is. During minimization and equilibration simulations, restraining potentials were applied to some lipid atoms to prevent integration from becoming unstable and to prevent the bilayer from deforming from non-optimal initial configurations. The strengths of the restraining potentials and the atoms to which the potentials were applied are listed in **Table S2**. Energy minimization used a steepest descent algorithm for up to 5000 steps or until machine precision was reached. The first two steps of equilibration were conducted in the Canonical ensemble (NVT) for 125 ps each with an integration timestep of 1 fs. Between these steps, the restraining potentials were relaxed. The third step of equilibration was in the isothermal-isobaric ensemble (NPT) and was conducted for 125 ps with an integration timestep of 1 fs.

Restraining potentials were further relaxed during this step. The C-rescale barsostat was turned on during the third step and remained turned on for the rest of the equilibration. In both the fourth and fifth steps, the restraining potentials are further relaxed and 250 to 500 ps of simulation were conducted in the NPT ensemble with an integration timestep of 2 fs. In the sixth step, restraints were removed and 500 ps of simulation are conducted in the NPT ensemble with an integration timestep of 2 fs. After equilibration, the box was inspected to determine whether the distance between periodic images of the lipid bilayer was at least 4.0 nm in the z direction. This distance ensured there were no artifacts from system size. Subsequent NPT simulations were used for data production.

**MD Data Production and Defect Analysis.** MD production simulations were conducted for 400 ns. Snapshots containing atomic coordinates of lipids were collected every 10 ps. The first 100 ns of data were excluded from analysis. Exclusion of the first 100 ns of data was meant to ensure that box volume and dimensions had converged to equilibrium values and to decouple the system coordinates from the initial structures produced by CHARMM-GUI or Packmol. Membrane defects were analyzed using the PackMem package [10]. As recommended by the PackMem authors, three thousand evenly spaced simulation snapshots were extracted from the last 300 ns of the simulation trajectories (1 frame per 100 ps). PackMem was used to identify shallow, deep, and all defects within snapshots.

The PackMem software analyzes a top-down, two-dimensional representation of the lipid membrane, detecting areas where aliphatic atoms are exposed. Specifically, PackMem generates a 1 Å by 1 Å grid. Each grid cell is examined to determine whether the atom closest to the membrane surface is an aliphatic atom. If an aliphatic atom is detected before any other atoms, the cell is considered to have a packing defect. The depth of a defect is determined by the vertical distance between the aliphatic atom and the central glycerol atom on the given lipid chain [10]. This vertical distance is compared to a threshold parameter, d. Any defect with a vertical distance less than or equal to d is considered a shallow defect. Any defect with a distance larger than d is considered a deep defect. In our analysis d was set to 1 Å. This distance has been shown to agree well with experimental observations [10]. Because cholesterol differs in structure from a phospholipid, a central glycerol atom is not present. Cholesterol's hydroxyl oxygen was used as a proxy to mimic the same analysis. The hydroxyl oxygen was chosen because it sits at nearly the same depth in the membrane as the surrounding lipid glycerol groups.

Defects were quantified with three different descriptors, namely percent defect coverage, defect size, and raw counts per frame. To examine the convergence of the defect descriptors, error bars are calculated by examining values calculated from the first third, middle third, and final third of the 3000 analyzed simulation frames. The sample standard deviation of these values is used to approximate the error as prescribed by the PackMem developers. Low error suggests that the simulation has reached an equilibrium state and that the membrane is no longer structurally evolving. The average defect coverage of the lipid membrane surface is calculated. This is done using the expected value for raw number of defects per frame and the expected size of defects (**Equation S1**).

$$\%A_{covered,avg} = 100\% * \frac{E(A_{defect})E(C)}{A_{lipid}} = 100\% * \frac{\sum_{A \in Areas} p_A A \sum_{C \in Counts} p_C C}{A_{lipid}} \qquad \text{(Eq. S1)}$$

$P_A$ denotes the probability any given defect will be of size A, and $P_C$ denotes the probability a given frame will have C counts. $E(A_{defect})$ and E(C) represent the expected value for the size and number of defects per frame, respectively. These are used in conjunction with the computed average membrane surface area, $A_{lipid}$, to calculate the expected percentage of the membrane surface consisting of defects. The membrane surface area fluctuates by less than 1% after reaching equilibrium, thus this contribution to the error is not considered. Percentage defect coverage allows a straightforward comparison between membranes in terms of area accessible to the α-synuclein. As recommended by the PackMem developers, defects with an area less than 15 $Å^2$ are excluded from analysis. Analysis of these small defects tends to be noisy, and convergence of values is difficult to attain. Because our analysis relies on raw defect counts rather than the fit distribution, all defects at or above the 15 $Å^2$ threshold are included. We present the "all" defects descriptor from PackMem which includes defects no matter how deep they are in the membrane. Experimentally observed increases in binding correlated with increases in the population and size of all defects in simulations. Simulated defect depth was not an explanatory factor for changes in experimentally measured binding.

**Volume Labeling Assay Methods and Discussion**

To ensure the imaged structures were unilamellar vesicles, a volume labeling assay was used **(Figure S9)**. SUVs containing lipid-anchored ATTO-647N were formed in the presence of 100 μM ATTO-488 dye, encapsulating it within the vesicle lumen. The vesicles were then tethered onto the glass surface as described in **Slide Passivation and Tethered Vesicle Assay**. All excess ATTO-488 dye in solution was rinsed away from outside the vesicles in the final buffer rinsing step. All vesicle compositions were then imaged under the same two-channel confocal settings with ATTO-488 representing the vesicle lumen (i.e. volume label) and ATTO-647N representing the vesicle bilayer (i.e., surface label). The following fluorescence intensity scaling behavior must be true for hollow spheres (i.e., unilamellar vesicles): $(lumen\ intensity\ or\ ATTO488) \sim diameter^3$ and $(membrane\ intensity\ or\ ATTO647N) \sim diameter^2$. Therefore, when the cube root of ATTO488 intensity is plotted vs the square root of ATTO647N intensity, the plot must exhibit linearity. Linear correlation was indeed observed between the two quantities, indicating the presence of hollow spheres or unilamellar vesicles.

**Poly-L-Lysine (PLL) Adsorption Assay Methods and Discussion**

Because DLPS has non-negligible water solubility with an estimated CMC of ~1 μM [12], we performed a tethered vesicle assay with poly-L-lysine (PLL) to verify DLPS was not leaching from the DLPC/DLPS vesicles and influencing the vesicle surface charge (**Figure S10**). PLL was labeled with ATTO 488 by incubating PLL with ATTO 488 NHS-Ester in a 1:1 molar ratio in 20mM HEPES, 150mM NaCl (pH 7.4) at room temperature for 30 minutes. Excess dye was removed by running the solution through a 7k MWCO Zeba Spin Column. The vesicles were prepared and imaged as described in the **SUV Preparation** and **Slide Passivation and Tethered Vesicle Assay**. 500 nM of PLL-ATTO488 was incubated with DOPC, 3:1 DOPC/DOPS, DLPC, and 3:1 DLPC/DLPS, and the fluorescence intensity of the PLL-ATTO488 adsorbed to 100 nm average vesicles was extracted. There was negligible adsorption to the membranes containing only PC lipids and strong adsorption to membranes containing PS lipids. Furthermore, the level of adsorption was comparable between membranes containing DLPS and DOPS, indicating that DLPS leaching was negligible in the tethered vesicle assay used.

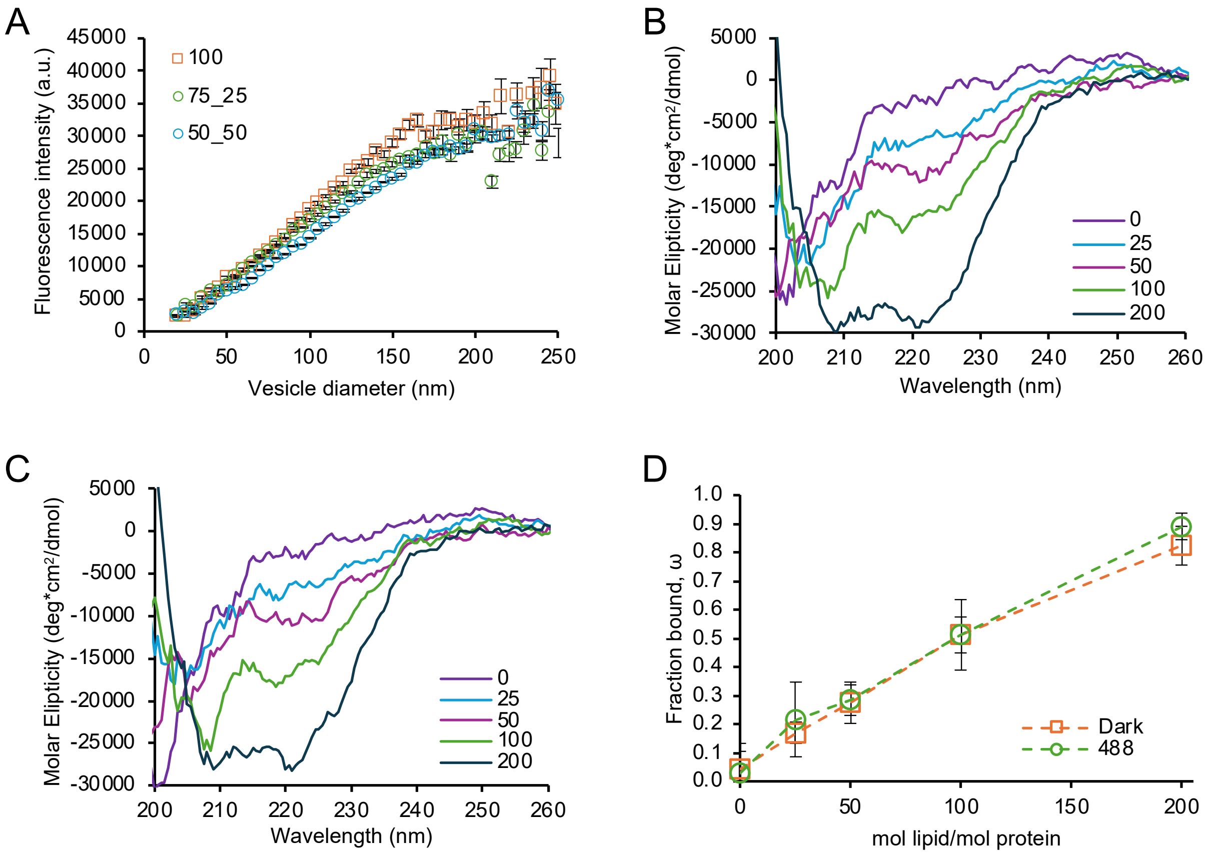


**Figure S1. Fluorescence Microscopy and CD spectroscopy reveal that ATTO 488 does not alter αSyn's binding affinity for membranes.** (A) Results from fluorescence microscopy-based tethered vesicle assay. Protein fluorescence intensity is plotted as a function of vesicle diameter on SUVs composed of 3:1 DPhPC:DPhPS. NTA-αSyn was incubated with vesicles at a total protein concentration of 250nM. Among 3 separate experiments, the ratio of labeled:dark NTA-αSyn was varied from 1:0, 3:1, and 1:1. In the experiments containing dark αSyn, the protein intensities were corrected by appropriate factors (i.e., either 75% or 50%) to account for the dark αSyn population. The 3 intensity profiles converge to comparable values, indicating that binding was not affected by fluorescent labeling with ATTO 488. Each data point represents a 5nm bin containing N = 17-374 vesicles. Error bars represent the standard error of the mean (SEM). (B-C) Results from CD spectroscopy using SUVs containing 3:1 DPhPC:DPhPS. (B) CD spectra for 1 μM NTA-αSyn-ATTO488 incubated with SUVs ranging from 0-200 μM lipid concentration. (C) CD spectra for 1 μM dark NTA-αSyn incubated with SUVs ranging from 0-200 μM lipid concentration. (D) The fraction of NTA-αSyn bound, ω, as a function of lipid concentration. The CD spectroscopy-derived binding curves for dark and fluorescent NTA-αSyn in panel D are nearly identical.

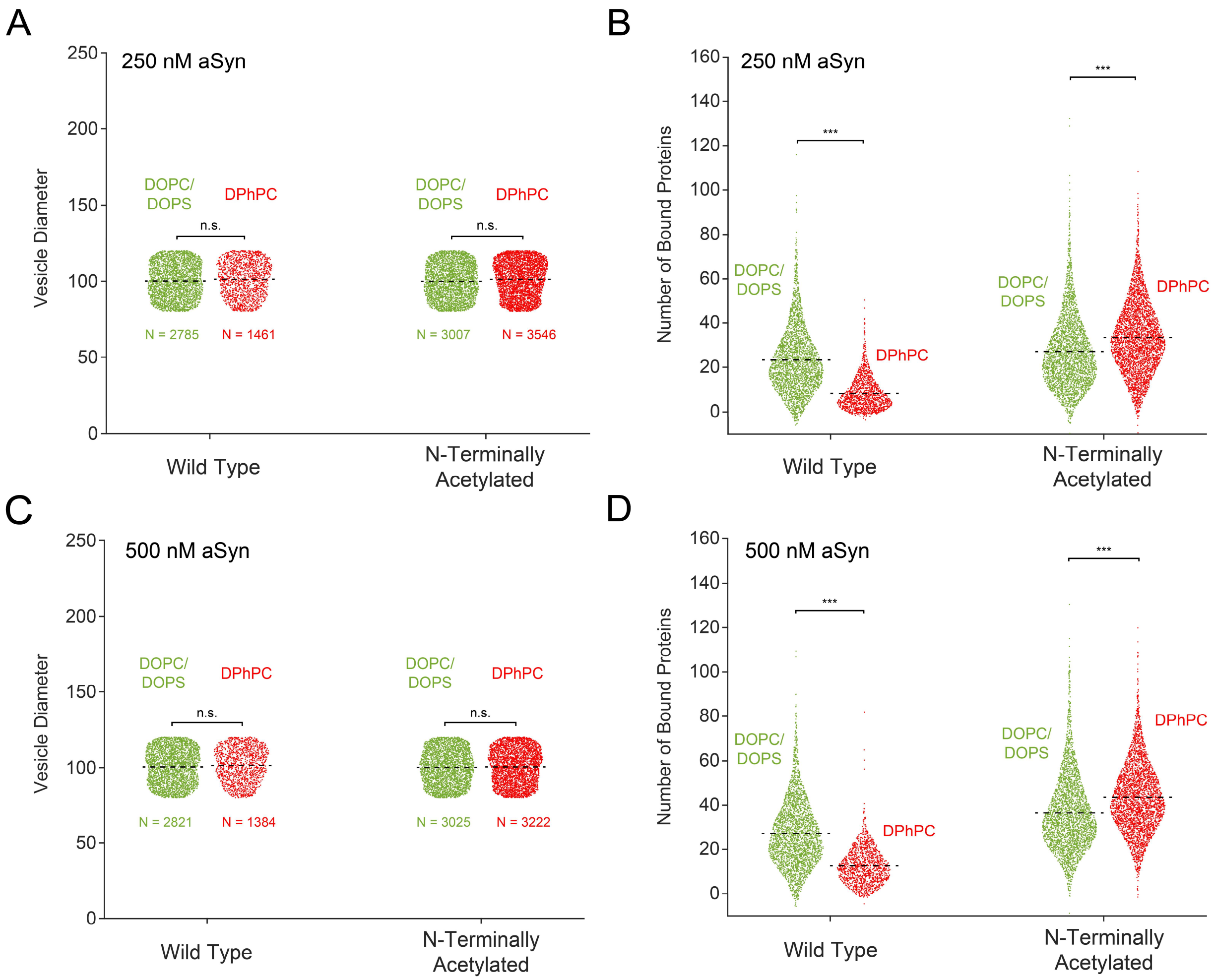


**Figure S2. Supplementary data for experiments comparing WT-αSyn and NTA-αSyn binding to DOPC/DOPS and DPhPC membranes.** (A) The distribution of diameters for analyzed vesicles that were incubated with 250 nM αSyn. (B) The number of proteins bound to vesicles in B. (C) The distribution of diameters for analyzed vesicles that were incubated with 500 nM αSyn. (n.s. = not significant as determined by unpaired Student's t-test). (D) The number of proteins bound to vesicles in C. Errors in A and C represent standard deviations. n.s. = not significant as determined by unpaired Student's t-test. *** = $P < 0.0001$ as determined by Welch's t-test (WT) and unpaired Student's t-test (NTA).

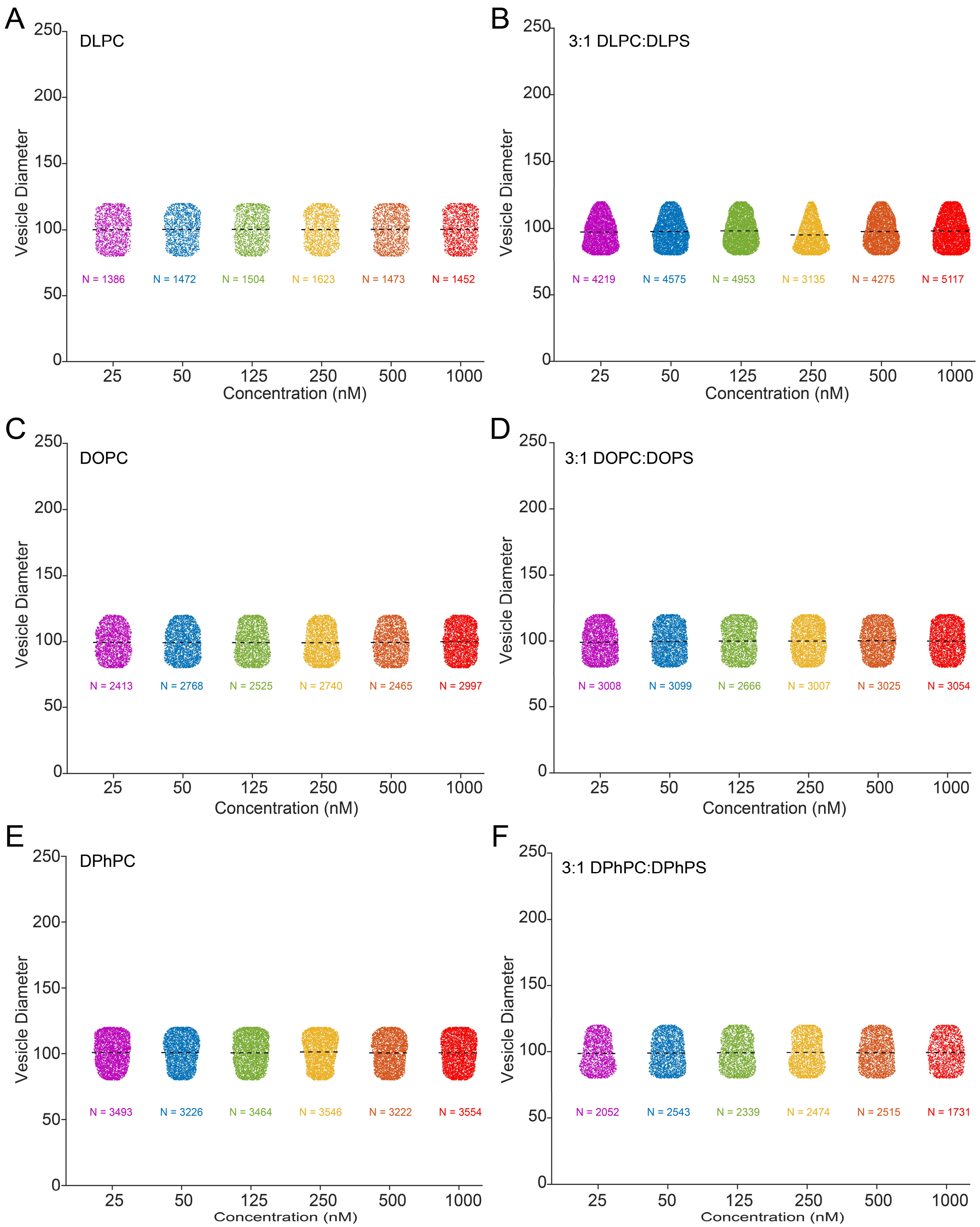


**Figure S3. Distributions of diameters for analyzed vesicles from Figures 3 and 4 of the main text.** Black dashed lines indicate the average diameter of the populations. All data sets possessed an average vesicle diameter of approximately 100 nm.

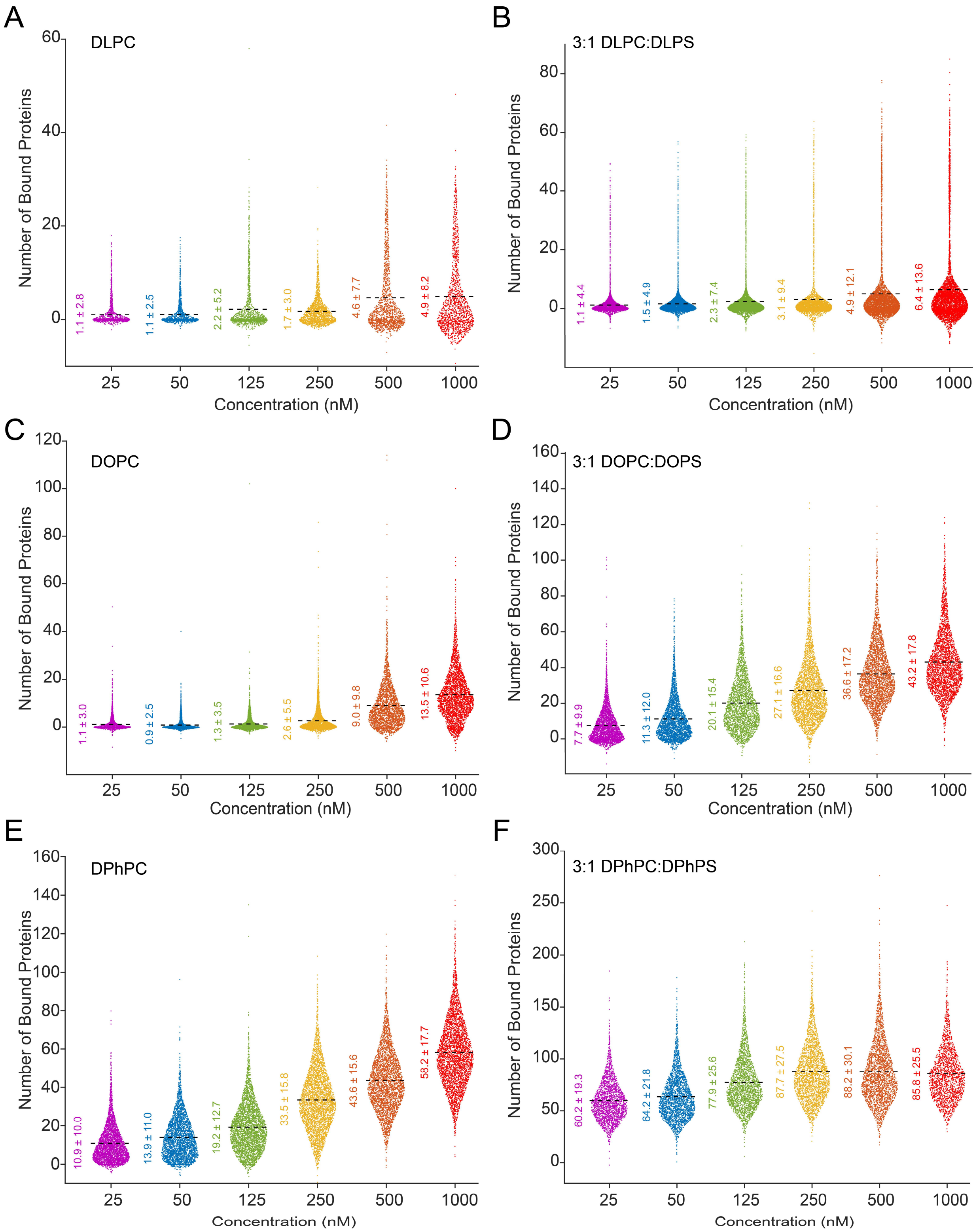


**Figure S4. Protein content on analyzed vesicles from Figures 3 and 4 of the main text.** The distribution of proteins bound to individual vesicles. These distributions are coupled to the distributions in Figure S3. The black dashed line and values indicate the average number of proteins bound. Errors listed represent standard deviations.

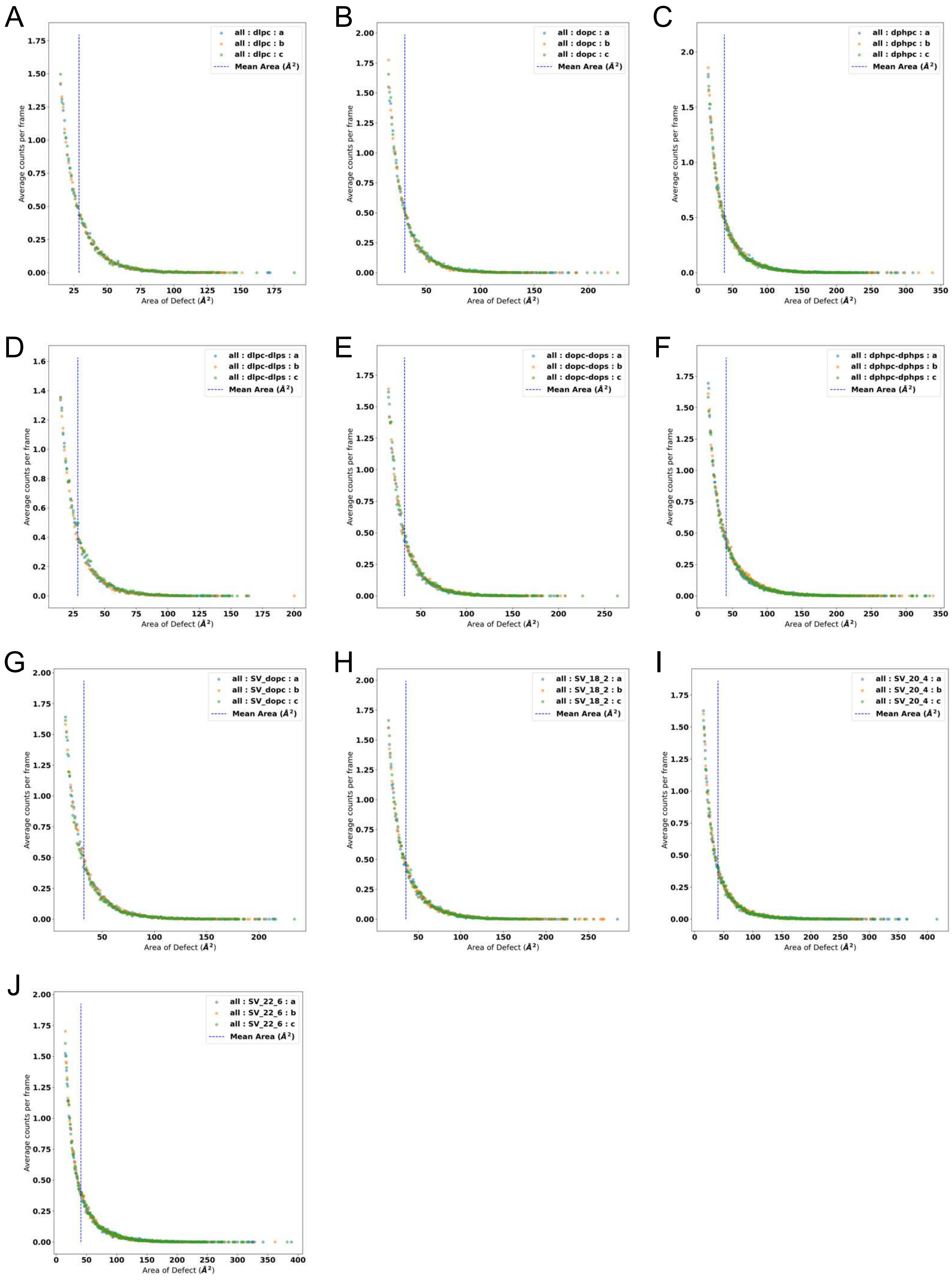
A
all : dlpc : a
all : dlpc : b
all : dlpc : c
Mean Area (Å²)
Average counts per frame
Area of Defect (Å²)
B
all : dopc : a
all : dopc : b
all : dopc : c
Mean Area (Å²)
Average counts per frame
Area of Defect (Å²)
C
all : dphpc : a
all : dphpc : b
all : dphpc : c
Mean Area (Å²)
Average counts per frame
Area of Defect (Å²)
D
all : dlpc-dlps : a
all : dlpc-dlps : b
all : dlpc-dlps : c
Mean Area (Å²)
Average counts per frame
Area of Defect (Å²)
E
all : dopc-dops : a
all : dopc-dops : b
all : dopc-dops : c
Mean Area (Å²)
Average counts per frame
Area of Defect (Å²)
F
all : dphpc-dphps : a
all : dphpc-dphps : b
all : dphpc-dphps : c
Mean Area (Å²)
Average counts per frame
Area of Defect (Å²)
G
all : SV_dopc : a
all : SV_dopc : b
all : SV_dopc : c
Mean Area (Å²)
Average counts per frame
Area of Defect (Å²)
H
all : SV_18_2 : a
all : SV_18_2 : b
all : SV_18_2 : c
Mean Area (Å²)
Average counts per frame
Area of Defect (Å²)
I
all : SV_20_4 : a
all : SV_20_4 : b
all : SV_20_4 : c
Mean Area (Å²)
Average counts per frame
Area of Defect (Å²)
J
all : SV_22_6 : a
all : SV_22_6 : b
all : SV_22_6 : c
Mean Area (Å²)
Average counts per frame
Area of Defect (Å²)

**Figure S5. Frequency distributions for defects on MD-simulated membranes.** Distributions for (A) DLPC, (B) DOPC, (C) DPhPC, (D) 3:1 DLPC:DLPS, (E) 3:1 DOPC:DOPS, (F) 3:1 DPhPC:DPhPS, (G) SV mimics containing DOPC, (H) SV mimics containing 18:2 PC, (I) SV mimics containing 20:4 PC, and (J) SV mimics containing 22:6 PC. Each of the 3 distributions in each panel is normalized by the total number of simulation frames – 1000 frames.

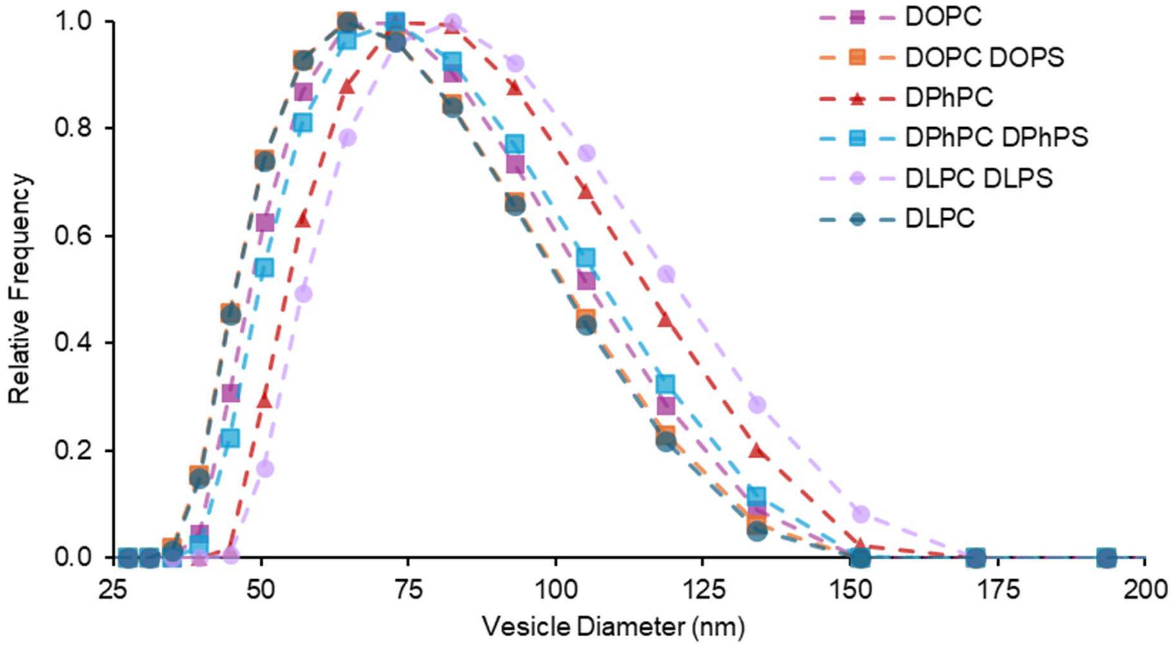


**Figure S6. DLS distributions of 50 nm extruded vesicles used for CD Spectroscopy.** The distributions shown are the average of 3 measurements for each sample.

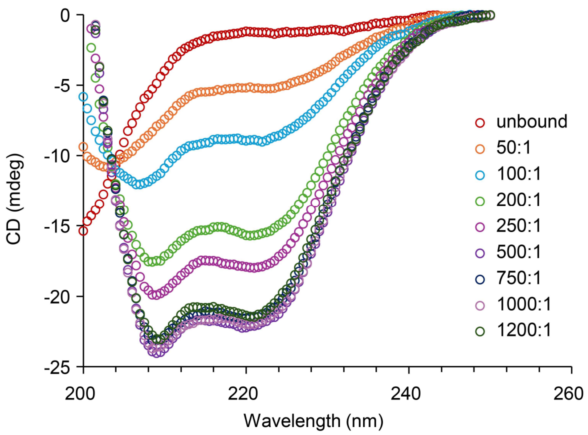


**Figure S7**. Circular dichroism spectra of NTA-αSyn for 3:1 DPhPC:DPhPS.

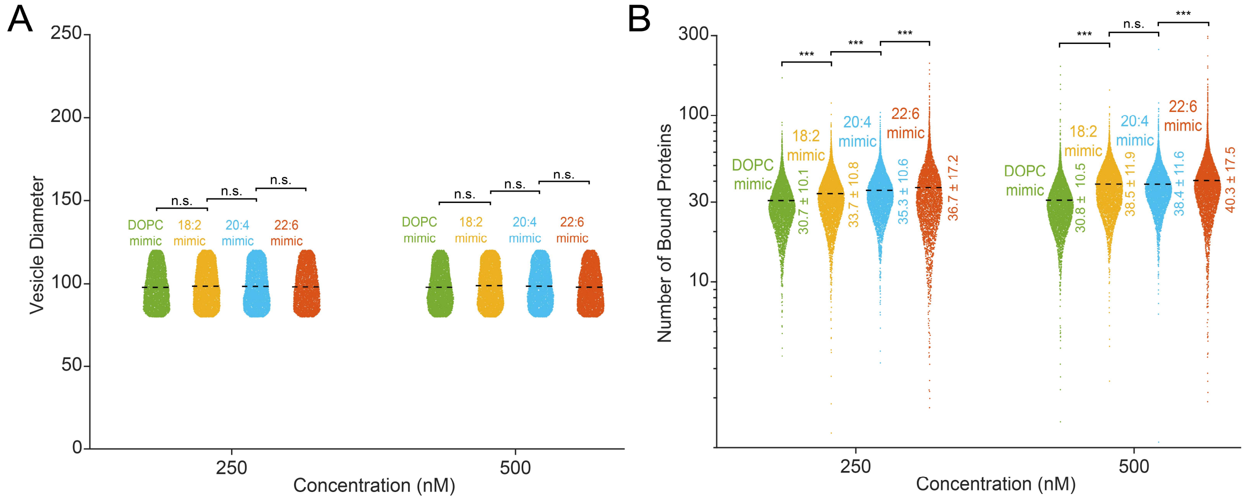


**Figure S8. Supplemental NTA-αSyn binding data for the SV mimics.** (A) The distributions of diameters for analyzed vesicles, which all possessed an average diameter of approximately 100 nm. The number of vesicles (N) analyzed within each condition are as follows: DOPC mimic with 250 nM αSyn (N = 4756), 18:2 PC mimic with 250 nM αSyn (N = 5660), 20:4 PC mimic with 250 nM αSyn (N = 5078), 22:6 PC with 250 nM αSyn (N = 4050), DOPC mimic with 500 nM αSyn (N = 5184), 18:2 PC mimic with 500 nM αSyn (N = 5431), 20:4 PC mimic with 500 nM αSyn (N = 4896), and 22:6 PC with 500 nM αSyn (N = 5047). (B) Number of proteins bound to vesicles in C that were incubated with 250 or 500 nM NTA-αSyn. The black dashed line and values on jitter plots in C and D indicate the average vesicle diameter and number of proteins bound, respectively. n.s. = not significant as determined by unpaired Student's t-test. *** = P < 0.001 as determined by unpaired Student's t-test.

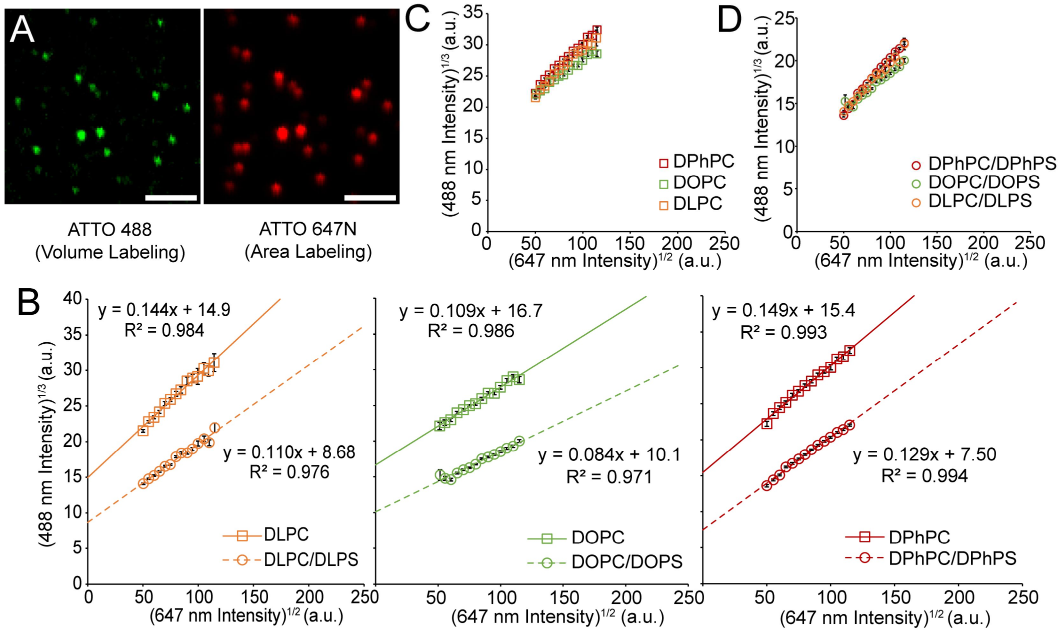


**Figure S9. Volume labeling assay.** (A) Fluorescence micrographs of SUVs encapsulating ATTO 488 dye for volume labeling and containing ATTO 647N in the bilayer for area labeling. Scale bars represent 2 μm. (B) Scaling relationship between SUV volume and surface area for PC membranes and 3:1 PC:PS membranes. The intensity values correspond to SUVs with an average diameter of 100 nm. (C) Overlay of the cube root of 488 nm intensity versus the square root of 647 nm intensity for DPhPC, DOPC, and DLPC. (D) Overlay of the cube root of 488 nm intensity versus the square root of 647 nm intensity for DPhPC/DPhPS, DOPC/DOPS, and DLPC/DLPS.

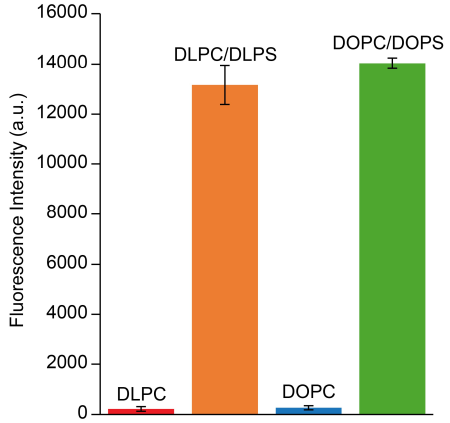


**Figure S10. Vesicle surface charge assay.** The average fluorescence intensity of ATTO 488-labeled PLL adsorbed to DLPC, 3:1 DLPC:DLPS, DOPC, and 3:1 DOPC:DOPS SUVs with diameters ranging from 90-110 nm.

**Table S1. Molecular membrane compositions for all bilayer simulations.**

| System | Composition (number of molecules) |
|---|---|
| *DPHPC* | 288 DPhPC |
| *DPHPC/PS* | 216 DPhPC, 72 DPhPS |
| *DLPC* | 288 DLPC |
| *DLPC/PS* | 216 DLPC, 72 DLPS |
| *DOPC* | 288 DOPC |
| *DOPC/PS* | 216 DOPC, 72 DOPS |
| *SV DOPC* | 100 DOPC, 100 DOPE, 40 DOPS, 160 Cholesterol |
| *SV 18:2* | 100 D(18:2)PC, 100 DOPE, 40 DOPS, 160 Cholesterol |
| *SV 20:4* | 100 D(20:4)PC, 100 DOPE, 40 DOPS, 160 Cholesterol |
| *SV 22:6* | 100 D(22:6)PC, 100 DOPE, 40 DOPS, 160 Cholesterol |

**Table S2. Details of the simulation equilibration and production steps.** These steps were repeated for each membrane composition.

| Step | Thermostat (K) | Barostat (bar) | Sim. Length (ps) | Timestep (fs) | Restraints | Strength (kJ/mol/nm) |
|---|---|---|---|---|---|---|
| *Equil. 1* | V-rescale, 303.15 | None | 125 | 1 | Head Groups, Dihedrals | 1000, 1000 |
| *Equil. 2* | V-rescale, 303.15 | None | 125 | 1 | Head Groups, Dihedrals | 400, 400 |
| *Equil. 3* | V-rescale, 303.15 | C-rescale, 1.0 | 125 | 1 | Head Groups, Dihedrals | 400, 200 |
| *Equil. 4* | V-rescale, 303.15 | C-rescale, 1.0 | 250 | 2 | Head Groups, Dihedrals | 200, 200 |
| *Equil. 5* | V-rescale, 303.15 | C-rescale, 1.0 | 500 | 2 | Head Groups, Dihedrals | 40, 100 |
| *Equil. 6* | V-rescale, 303.15 | C-rescale, 1.0 | 500 | 2 | None | N/A |
| *Production* | V-rescale, 303.15 | Parrinello-Rahman, 1.0 | 400,000 | 2 | None | N/A |

**Table S3. Calculated fraction bound (ω) of NTA-αSyn from CD Spectroscopy.** Each lipid composition and lipid:protein ratio tested is shown. The errors listed are the standard error of the mean.

| Fraction Bound (ω) | | | | | | | | |
|---|---|---|---|---|---|---|---|---|
| | mol lipid/mol protein | | | | | | | |
| Lipid Composition | 50:1 | 100:1 | 200:1 | 250:1 | 500:1 | 750:1 | 1000:1 | 1200:1 |
| DLPC | 0.010 ± 0.002 | 0.016 ± 0.002 | -0.005 ± 0.002 | 0.018 ± 0.002 | 0.001 ± 0.005 | -0.010 ± 0.004 | -0.010 ± 0.007 | 0.015 ± 0.004 |
| DOPC | 0.046 ± 0.006 | 0.048 ± 0.002 | 0.068 ± 0.003 | 0.029 ± 0.009 | 0.149 ± 0.010 | 0.178 ± 0.012 | 0.215 ± 0.017 | 0.295 ± 0.017 |
| DPhPC | 0.069 ± 0.004 | 0.150 ± 0.002 | 0.299 ± 0.002 | 0.367 ± 0.003 | 0.633 ± 0.003 | 0.795 ± 0.002 | 0.911 ± 0.002 | 0.926 ± 0.002 |
| DLPC/DLPS (3:1) | -0.010 ± 0.002 | 0.016 ± 0.001 | -0.003 ± 0.005 | 0.009 ± 0.002 | 0.033 ± 0.001 | 0.044 ± 0.002 | 0.037 ± 0.005 | 0.032 ± 0.005 |
| DOPC/DOPS (3:1) | 0.069 ± 0.004 | 0.114 ± 0.008 | 0.263 ± 0.005 | 0.328 ± 0.005 | 0.607 ± 0.003 | 0.759 ± 0.004 | 0.806 ± 0.011 | 0.814 ± 0.010 |
| DPhPC/DPhPS (3:1) | 0.175 ± 0.007 | 0.359 ± 0.005 | 0.691 ± 0.003 | 0.811 ± 0.002 | 1.024 ± 0.001 | 0.989 ± 0.002 | 1.015 ± 0.002 | 0.972 ± 0.002 |

**Table S4. Apparent dissociation constant ($K_d$) and cooperativity (n) determined by Langmuir-Hill fit to binding curves.** $K_d$ and cooperativity values were determined by fitting data in Figures 3E and 4B to Equation 2 from the main text. The errors listed are the standard error of the mean.

| Lipid Composition | $K_d$ | n |
|---|---|---|
| DOPC | 3376 ± 897 | 0.97 ± 0.17 |
| DPhPC | 330 ± 13 | 1.69 ± 0.10 |
| DOPC/DOPS | 386 ± 13 | 1.49 ± 0.07 |
| DPhPC/DPhPS | 125 ± 8 | 1.99 ± 0.21 |
